# Learning interpretable kinetic models for biomolecular interaction networks

**DOI:** 10.64898/2026.09.14.751382

**Authors:** Roi Eliasian, Yael Hazan, Timna Tzori, Barak Raveh

**Affiliations:** School of Computer Science and Engineering, The Hebrew University of Jerusalem, Jerusalem, Israel

**Keywords:** Molecular Dynamics, Markov State Model, Nuclear Pore Complex

## Abstract

Many cellular machines operate through weak, transient, and multivalent interactions whose functional states are governed by recurring interaction patterns rather than persistent molecular geometries. Here, we introduce interaction-based Markov state models (iMSMs), which construct interpretable kinetic models using unsupervised clustering of time-averaged, identity-resolved interaction distributions around a focal entity. Applied to nucleocytoplasmic transport at two molecular resolutions, iMSMs resolve graded interaction states spanning strong, partial, and weak engagement. During pore transport, partially engaged states provide faster routes to disengagement than strongly bound states, while the networks reveal transport pathways, interaction hubs, bottlenecks, and kinetic commitment. At a finer resolution, FG motifs exchange contacts while remaining associated, recovering established slide-and-exchange dynamics. iMSMs reproduce free energies, permeabilities, and multi-step kinetics, with nearly fourfold faster permeability convergence from truncated trajectories than direct transport-event counting. Thus, iMSMs connect rapidly exchanging contacts to graded interaction states and their kinetics across molecular resolutions.

## 1 Introduction

Many biomolecular processes are governed not by transitions between a small number of stable molecular structures, but by weak, transient, and multivalent interactions whose functional consequences emerge from their statistical organization over time[1–4]. This principle spans molecular scales: from fuzzy interactions of intrinsically disordered motifs with individual protein surfaces[1, 3, 5], to transport through disordered molecular barriers[6, 7], to mesoscale signaling assemblies and biomolecular condensates[2, 4, 8–10]. At larger scales, such systems can be viewed as dynamic biomolecular networks (DBNs), whose composition and connectivity continually reorganize rather than forming fixed, stoichiometrically defined complexes. DBNs include liquid-like biomolecular condensates such as stress granules, membrane-associated signaling clusters such as LAT signalosomes, and spatially anchored networks of intrinsically disordered domains, such as the FG repeats mediating nucleocytoplasmic transport through nuclear pore complexes (NPCs) [4, 7, 8, 10–13]. Despite substantial differences in composition, geometry, and scale, these systems pose a common computational challenge: extracting mechanistically interpretable kinetic models from high-dimensional simulation trajectories. Such models should identify recurring interaction environments, retain their molecular meaning, and describe how their interconversion relates to function.

Among these systems, nucleocytoplasmic transport through NPCs provides a particularly informative example. The NPC is the main gateway for regulated molecular exchange between the nucleus and the cytoplasm, and it therefore plays an essential role in gene expression, RNA export, signaling, mechanical transduction, aging, and disease, including cancer onset and viral infection[7, 14–20]. While its molecular composition and architecture are relatively well characterized, its function emerges from a dynamic network of transient interactions. Its massive, eightfold symmetric ring scaffold is composed of structured domains from some 500 nucleoporin protein subunits (Nups). This scaffold surrounds a central channel occupied by intrinsically disordered phenylalanine-glycine (FG)-repeat domains from a subset of nucleoporins termed FG Nups [21–24]. The FG Nups vary in sequence composition, FG-motif distribution, and chain length, and are anchored at distinct positions throughout the NPC scaffold, creating a chemically and spatially heterogeneous transport barrier [6, 22, 25]. They selectively block the passive diffusion of large inert macromolecules while facilitating the rapid transport of cargo-bound nuclear transport receptors (NTRs) [7, 26–30]. Because the central channel is densely populated by FG-repeat domains from multiple FG Nup chains, a cargo:NTR complex may encounter a changing, multivalent interaction environment during translocation. Within this highly dynamic environment, its pathway and kinetics may be influenced by cargo or NTR size, receptor valency, FG-repeat composition and spatial organization, pore geometry, and the Ran-driven transport cycle [6, 7, 31, 32].

Experimental and theoretical advances have clarified many aspects of NPC architecture and transport. Integrative studies mapped yeast and human NPC scaffolds [21–23]; in situ cryo-electron tomography showed that the cellular environment shapes NPC architecture [24]; reconstituted FG-repeat systems and artificial pores tested whether simplified barriers reproduce selectivity [29, 31]; single-molecule, AFM, and super-resolution approaches probed barrier organization and dynamics [32–34]; and physical modeling linked polymer organization, transport-receptor binding, and permeability [6, 7]. Together, these studies suggest that selective transport emerges from fuzzy, multivalent and spatially organized NTR–FG-repeat interactions. However, important questions remain only partly resolved: How are FG-repeat domains functionally arranged along transport routes? Do cargoes use a common central path, multiple peripheral conduits, or choose routes depending on receptor properties? How do cargo size, receptor valency, and FG-repeat motif organization affect dwell time, lateral mixing between spokes, arrest probability, and bottlenecks? How do geometric or mechanical pore perturbations alter local cargo–barrier encounters [6, 7, 20, 34]?

Answering these questions requires methods that retain information about which components interact and how those interactions change over time, while compressing the underlying trajectories into an interpretable kinetic representation. Molecular dynamics (MD) simulations provide time-resolved, molecular-level views of component motions and interactions, while advances in coarse graining, enhanced sampling, machine-learned potentials, and high-performance computing enable their application to increasingly large systems[35–40]. Nevertheless, accessible system sizes and simulation times remain limited, particularly when transitions between relevant states are rare[35, 40, 41]. Converting these trajectories into compact mechanistic models remains a separate challenge. Predicting observables or compressing trajectories is not sufficient for this purpose: the inferred states must admit a molecular interpretation, and their interconversion must retain the relevant kinetics. In highly dynamic systems, a functional state may encompass many microscopic configurations rather than a single conformation. The challenge is therefore one of representation as well as sampling: the system’s relevant state may be better described by a recurring statistical pattern of interactions among different types of components, rather than by the instantaneous coordinates of individual molecules or the exact set of pairwise contacts between specific molecular copies.

Markov state models (MSMs) provide a general framework for extracting long-timescale kinetics from molecular simulations by partitioning trajectory data into discrete states and estimating transition probabilities between them [41–43]. MSMs can summarize complex trajectories, estimate stationary and kinetic observables, and infer long-timescale behavior from sets of shorter simulations. Their scope has expanded through unsupervised and deep-learning approaches to kinetic model construction, alongside multiscale and decomposed representations [44–48]. As MSMs have been applied to increasingly complex molecular processes, considerable effort has gone into choosing state representations that capture the relevant slow dynamics without retaining unnecessary structural detail. In molecular recognition, for example, protein–ligand contacts have been used directly as features for MSM construction [49, 50]; more recently, intermolecular contact maps combined with structural descriptors have resolved encounter complexes and partially bound fuzzy states during binding of an intrinsically disordered protein [51]. Other approaches have shifted the representation from global conformations toward local or temporally extended descriptions: graph dynamical networks learn Markovian representations of the instantaneous local environment around selected target atoms or molecules, while segment-based clustering methods incorporate temporal information by classifying distributions of configurations over trajectory segments [52, 53]. Related ideas have also been applied beyond conventional folding and binding problems: MSMs have been used to describe exchange of individual chains between condensed and dilute phases of biomolecular condensates [54], while dynamic graphical models, independent Markov decomposition, and iVAMPnets reduce the combinatorial complexity of large systems by representing their dynamics in terms of local or weakly coupled subsystems [48, 55, 56]. Other extensions couple MSMs to reaction–diffusion dynamics [57]. Self-assembly MSMs use bond-based state representations to resolve assembly pathways [58, 59] and support optimization of time-dependent assembly protocols [60].

Together, these developments establish that molecular interactions, local molecular environments, and finite-time trajectory segments can all provide informative kinetic representations. For dynamic biomolecular networks governed by fuzzy and multivalent interactions, such a representation is particularly natural: individual contacts may form and break rapidly, and many distinct microscopic configurations may nevertheless correspond to the same persistent statistical pattern of engagement. Such dynamics may therefore be more naturally described by graded interaction states spanning strong, partial, and weak engagement than by sharply separated bound and unbound configurations.

Here, we introduce interaction-based Markov state models (iMSMs), which combine a physically specified interaction representation with unsupervised state identification to construct interpretable kinetic models. Each time window is represented by the distribution of interactions around a focal entity, and recurring distributions define kinetic states. By coarse-graining over microscopic geometry while retaining interaction-partner identity and time-averaged engagement, iMSMs are particularly suited to systems in which function emerges from heterogeneous, graded, and rapidly exchanging contacts. In nucleocytoplasmic transport, for example, a NTR need not be described as encountering a sequence of well-defined complexes, but rather as experiencing a changing statistical mixture of interactions with multiple copies and types of intrinsically disordered FG nucleoporins. We apply iMSMs to coarse-grained Brownian dynamics simulations of transport through an experimentally informed yeast NPC model [7]. The resulting models recover key stationary and kinetic properties of the underlying simulations while providing interpretable maps of transport pathways across cargo sizes and NTR valencies, including graded engagement states that connect persistent binding to disengagement. We further apply the same abstraction at finer molecular resolution to an FG motif interacting with Kap95, where the states resolve residue-level contact exchange during sliding. Thus, the same interaction-centered representation can describe recurring kinetic environments across distinct molecular resolutions, from local fuzzy binding to transport through a heterogeneous network of disordered proteins.

## 2 Results

### 2.1 Kinetic modeling via interaction-based Markov State Models (iMSMs)

iMSMs automate the identification of recurring interaction states and estimation of their transition network within a user-specified physical representation. By discretizing time-averaged interaction distributions, iMSMs coarse-grain over many microscopic contact configurations that need not share the same local geometry, provided that they preserve the same statistical interaction pattern around a selected focal entity. For example, an iMSM can describe how the interaction environment of a cargo:NTR complex changes as it traverses the central channel of the NPC. In general, the approach is applicable when the dynamics of interest can be represented by recurring local interaction environments that are approximately Markovian at an appropriate lag time.

An iMSM is constructed in five steps (Fig. 1a). First, given a set of molecular dynamics (MD) trajectories, the simulated system is partitioned into a set of discrete interacting components, *C* = {*c*_1_*, … , c_N_*}, such as the different FG Nups, together with a focal entity *f*, such as a cargo:NTR complex, that may interact with subsets of these components. Second, each MD trajectory is converted into an interaction trajectory *I*(*t*) that records the contacts of *f* with the components in *C*. Third, *I*(*t*) is divided into consecutive time windows of duration *τ* ; the interaction statistics in each time window are summarized in an interaction histogram *H*, which captures both the identity of the interacting components and the degree of engagement of the focal entity over time. The histogram is normalized by a fixed reference interaction capacity for the focal entity, rather than by the number of interactions actually observed, allowing the representation to distinguish strongly, partially, and weakly engaged states even when they involve similar interaction partners. This interaction-based feature representation can be augmented with process-specific observables when they improve the description of the relevant dynamics. Fourth, unsupervised clustering of the histograms identifies a reduced set of recurring interaction states, *S*= {*s*_1_*, … , s_k_*}. Fifth, transitions between these states are recorded and used to infer a lagged transition-probability matrix,

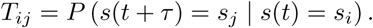

**Fig. 1.**
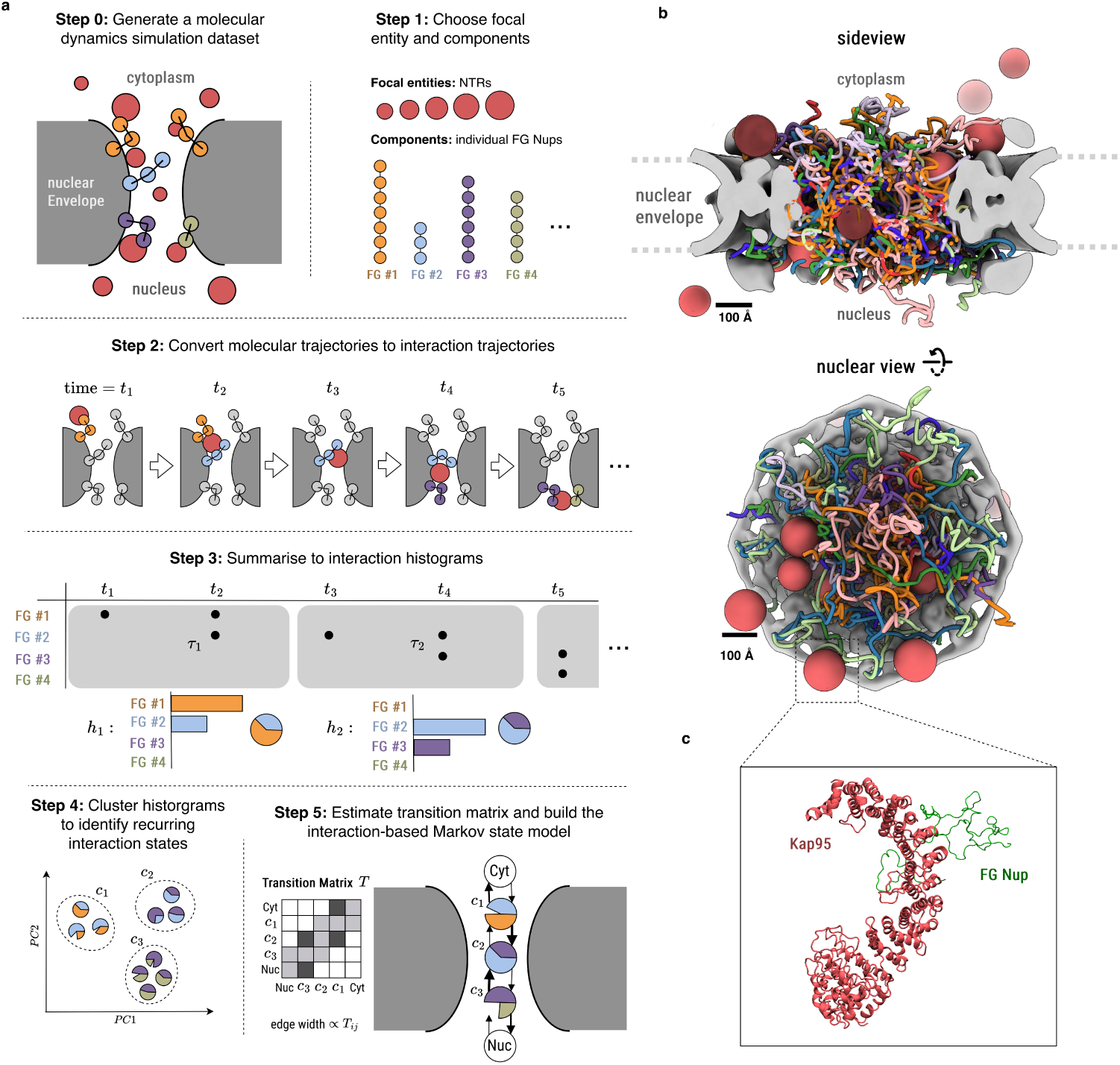
Unsupervised construction of an interpretable interaction-based kinetic model for nucleocytoplasmic transport. **(A)** A schematic overview of the iMSM workflow. Molecular simulation trajectories are generated, and a focal entity-here, an NTR, and its interacting components are selected. Coordinate trajectories are converted into time-dependent interaction trajectories, which are divided into windows of duration *τ* and summarized as normalized interaction histograms. Within this specified representation, unsupervised clustering identifies recurring interaction states, and observed transitions between states are used to estimate the transition-probability matrix *T* and construct the iMSM. Node sectors show the interaction composition of each inferred state, arrows indicate transitions and edge widths are proportional to transition probabilities. Cyt and Nuc denote cytoplasmic and nuclear non-interacting states, respectively. **(B)** Representative side and nucleoplasmic views of the coarse-grained yeast nuclear pore complex simulation system. Disordered FG-repeat domains occupy the central channel, together with the transported NTRs. Pore diameter, 54 nm. Scale bars, 100 Å. **(C)** Residue-level simulation system used for the molecular-scale application, showing Kap95 (red) interacting with an Nsp1-FSFG125 construct containing six FSFG motifs (green).

This matrix provides a compact kinetic model from which stationary distributions, transport observables and pathway-level quantities are estimated. The focal entity, interacting components, contact definition, window duration and clustering resolution are modeling choices, whereas state assignments and transition probabilities are inferred from the trajectories. Detailed choices for the interacting components, focal entity, feature construction, and probability estimation are provided in the Supplementary Methods.

### 2.2 iMSMs for studying facilitated nucleocytoplasmic transport

To demonstrate the iMSM framework, we constructed iMSMs to describe facilitated nucleocytoplasmic transport through the nuclear pore complex (NPC), where the cargo-bound NTRs cross the selective barrier by transiently interacting with the disordered FG repeat domains occupying the central channel[22, 27, 61]. Because this challenging process is governed by many transient NTR–FG-Nup contacts, it provides a natural testbed for interaction-based kinetic modeling: the goal is to compress large transport simulations into interpretable state models while preserving key properties of the underlying thermodynamic ensemble and transport kinetics, including state probabilities and transition rates.

We generated the input trajectories using the recent integrative coarse-grained Brownian dynamics model of nucleocytoplasmic transport through the yeast NPC, which was previously constrained and evaluated against diverse experimental measurements[7] (Fig. 1b). We then construct iMSMs from these trajectories and assess how well they reproduce thermodynamic and kinetic properties obtained by direct analysis of the corresponding full-length MD trajectories.

Specifically, for each of the 216 FG Nup chains, spanning ten FG Nup types[22], the intrinsically disordered FG-repeat domain was divided into N- and C-terminal components, thereby distinguishing regions proximal and distal to the structured anchoring domain that tethers the FG Nup to the NPC scaffold. We defined the cargo-bound NTRs as the focal entities whose evolving interaction environment we model. For each trajectory, we recorded the time-dependent contacts between each focal entity and the FG-Nup components. All NTR variants were represented using the same fixed interaction capacity of five. At each time point, the fraction of this capacity occupied by FG-Nup contacts was assigned to the corresponding FG-Nup components, while any remaining capacity was assigned to one of five spatially defined unoccupied categories according to the NTR position: the nucleus, cytoplasm, or one of three layers of the central channel. Contacts were defined as a surface-to-surface distance of ≤ 1 nm. The resulting interaction trajectories were partitioned into consecutive 5 *µ*s windows, with the window duration chosen after examining lag-dependent implied timescales (Supp. Fig. S1) [43, 62]. Each window was then represented by a normalized histogram of interaction frequencies. We then clustered the histograms to find recurring interaction environments, with the mean *z*-coordinate of the NTR within each window as an additional clustering feature. We used a common initial clustering resolution of 320 clusters for all NPC systems, followed by identical state-consolidation and minimum-occupancy criteria. This fixed resolution was used throughout rather than being optimized separately for individual NTR or pore conditions; full clustering and transition-estimation details are provided in the Supplementary Methods. To increase the effective sample size, we also utilized the eightfold rotational symmetry of the NPC scaffold, adjusting the interaction trajectory generation and histogram clustering steps to account for this symmetry while ensuring that symmetry-equivalent transport environments were represented consistently (Fig. 1b, Supp. Fig. S2).

We begin by inspecting two representative iMSMs, generated from transport trajectories of 61-kDa NTRs with either 4 or 6 FG-Nup interaction sites (Fig. 2a; Supp. Fig. S3). For non-interacting molecules of the same molecular weight, passive diffusion is orders of magnitude slower[7]; transport of these NTRs is therefore dominated by interactions with FG repeats. Both iMSMs closely reproduce key properties of the system’s thermodynamic ensemble and transport kinetics. For each FG Nup, we estimated the stationary NTR-interaction probability from either the iMSM or directly from the MD trajectories, and converted these probabilities into relative interaction free energies by Boltzmann inversion[7]. For both the 4-site and 6-site NTRs, the resulting free energies inferred from the iMSM closely match those estimated directly from the MD trajectories (Fig. 2b; Jensen–Shannon divergences = 0.001/0.048; Wasserstein distances = 4.4/9.2 nm, respectively). The remaining discrepancies are small and arise mainly from a slight overestimation of the cytoplasmic and nucleoplasmic unbound states, which in turn, slightly reduces the inferred probabilities of the remaining states. Both iMSMs also recapitulate the transport permeability of the corresponding MD simulations: permeability estimates obtained from random walks over the states of each iMSM converge to those obtained by directly tallying transport events in the fulllength MD trajectories (Fig. 2c). To compare convergence, we repeatedly evaluated both estimates using progressively larger subsets of the available data. Convergence is similar whether the input data were reduced by using shorter trajectories (Fig. 2c, top) or by using fewer full-length trajectories (Fig. 2c, bottom).

**Fig. 2.**
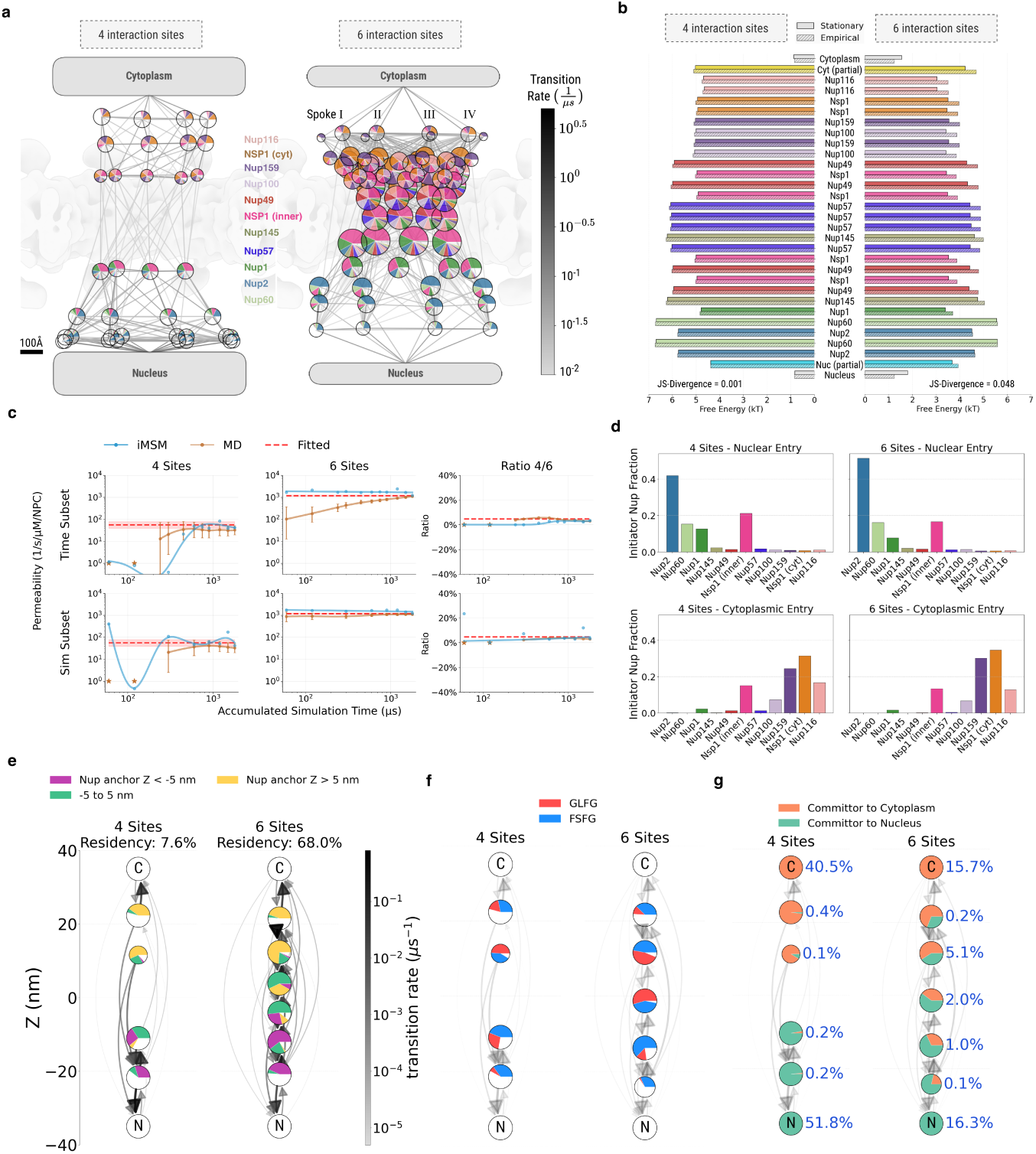
iMSMs reproduce transport properties and resolve valency-dependent NPC interaction pathways. **(A)** iMSMs for 61-kDa NTRs with four or six FG-Nup interaction sites in a 54-nm-diameter NPC. Nodes represent interaction states; colored wedges indicate FG-Nup contributions, white sectors represent unoccupied reference interaction capacity, and edges indicate transition rates computed using the infinitesimal generator matrix. State positions are estimated from the anchor positions of their constituent FG Nups. Cytoplasmic and nuclear unbound states are shown at the top and bottom, respectively. Node size is proportional to stationary probability. Scale bar, 100 Å. **(B)** Relative interaction free energies obtained from the iMSM stationary distribution or directly from MD trajectories. Jensen–Shannon divergences between the corresponding distributions are indicated. **(C)** Convergence of permeability estimates from iMSMs and direct MD transport counts as a function of accumulated simulation time, using either progressively longer trajectory segments (top) or increasing numbers of complete trajectories (bottom). The right panels show the four-site to six-site permeability ratio. Fit was determined through power law regression across MWs per site number. **(D)** Fractions of FG Nups initiating entry from the nuclear side (top) or cytoplasmic side (bottom), calculated from iMSM-generated transport trajectories. **(E-G)**, Single-spoke representations of the interaction states. States are colored by FG-Nup anchor position (E), GLFG or FSFG motif composition (F), or committor probability towards the cytoplasm or nucleus (G). Blue labels indicate stationary probabilities; residency denotes total stationary probability across all significantly non-empty interaction states.

#### 2.2.1 FG-Nup organization along transport pathways

The iMSMs provide compact descriptions of the transport pathways of the different representative NTRs and the role of different FG Nups in transport (Fig. 2a, Supp. movie 1). Across FG Nups and regardless of NTR valency, interaction probabilities are closely correlated with the relative contributions to the total mass of FG repeats (Supp. Fig. S4). However, some Nups contributed slightly more to NTR interactions than their mass shares predicted. For example, the contribution of both inner- and outer-ring Nsp1 copies was ~ 1.2-fold higher than expected by their mass alone.

We next examined the states adjacent to the unbound cytoplasmic and nuclear states, which we term initiator states. These states enabled us to identify the corresponding initiator Nups that mediate the initial docking of free-diffusing NTRs to the NPC (Fig. 2a,d). As anticipated, on the cytoplasmic side, the initiator states are dominated by the outer-ring copies of Nsp1, Nup159, and, to a lesser extent, Nup116 and Nup100. Conversely, on the nucleoplasmic side, these states are dominated by basket Nups Nup1, Nup2, and Nup60, and when normalized by mass, also Nup145 (Fig. 2d). Nsp1 emerges as the dominant nucleoporin across the central channel. This prominence is consistent with its extended length, high stoichiometry, and distribution across multiple layers along the *z*-axis. Interestingly, the inner-ring copies of Nsp1 also contribute to the initiator states on both sides of the NPC, despite their anchoring at the central channel constriction [22], even when normalized by their considerable mass (Supp. Fig. S5), highlighting the highly dynamic nature of the FG-repeat domains and their ability to extend far from their anchoring sites to sample broad regions of space. This spatial reach is further illustrated by the single-spoke pathway representation in Fig. 2e.

FG Nups contain distinct FG-repeat classes, including GLFG and FSFG motifs. Whether this sequence heterogeneity simply tunes the overall material properties of the barrier, or instead gives rise to spatially distinct interaction environments with distinct functional roles during NTR transport, remains unresolved [7, 22, 30, 63–65]. Because iMSM states retain the identities of their constituent FG Nups, we can convert the states to FG-motif histograms by quantifying the relative contributions of GLFG- and FSFG-containing Nups (Fig. 2f). Through this, iMSMs map the spatial organization of NTR interactions with FG motifs along the NPC’s central axis. GLFG and FSFG contributions were approximately balanced in states near the pore center and toward the cytoplasmic side, whereas FSFG motifs contributed more strongly to states on the nucleoplasmic side. These results support the existence of distinct FG-motif interaction environments along the transport pathway, although whether these environments perform distinct functional roles remains to be determined.

Together, these analyses indicate that some transport properties are largely conserved between the 4-site and 6-site NTRs, including the identities of prominent initiator Nups and the spatial pattern of FG-motif usage. As expected, the higher-valency 6-site NTRs are nonetheless substantially more likely to occupy innerring-proximal states than the 4-site NTRs, nearer to the constriction of the central channel (Fig. 2a,b), also leading to the identification of a substantially wider range of unique interaction states.

On the nucleoplasmic side, Nup2 exhibits a marked dominance for 6-site NTRs, whereas in 4-site variants, it competes more evenly with Nsp1, Nup1, and Nup60 (Fig. 2a). These kinetic differences should be interpreted cautiously, however, because Nup2 is known to be more mobile than other basket nucleoporins and to dynamically associate with different regions of the pore in vivo [66].

The two NTRs also differed markedly in their committor profiles (Fig. 2g). For 4-site NTRs, states in either half of the central channel were already strongly committed toward exiting from the corresponding side of the NPC, producing a sharp committor inversion near the pore center. In contrast, 6-site NTRs retained substantial probabilities of exiting through either side of the pore for most in-channel states, while remaining biased toward the side on which they were located. These models allow us to examine not only where NTRs bind, but how transitions between interaction environments connect persistent association to disengagement.

### 2.3 Large-Scale Screening and Validation of Transport Dynamics

To evaluate the generalizability and robustness of the iMSM framework, we generated a comprehensive evaluation dataset comprising coarse-grained MD simulations of 15 distinct nuclear transport receptor variants. To systematically map how NTR biophysical properties affect transport-associated interaction dynamics, we simulated NTRs across five different molecular weights (3.5, 9.5, 20.3, 37.0, and 61.1 kDa) and three distinct interaction valencies (2, 4, and 6 FG-Nup binding sites). Each of these 15 NTR variants was simulated across four physiologically relevant pore diameters (46, 54, 62 and 70 nm), establishing a multi-dimensional transport dataset. For dilation studies, ring radii were scaled while maintaining scaffold stoichiometry (216 FG-Nup chains), thereby decreasing FG density with increasing diameter. For each NTR and pore variant, we constructed an independent iMSM from its transport trajectories, tracking the evolving interaction environment of each NTR variant as it traverses the NPC (Fig. 3a).

**Fig. 3.**
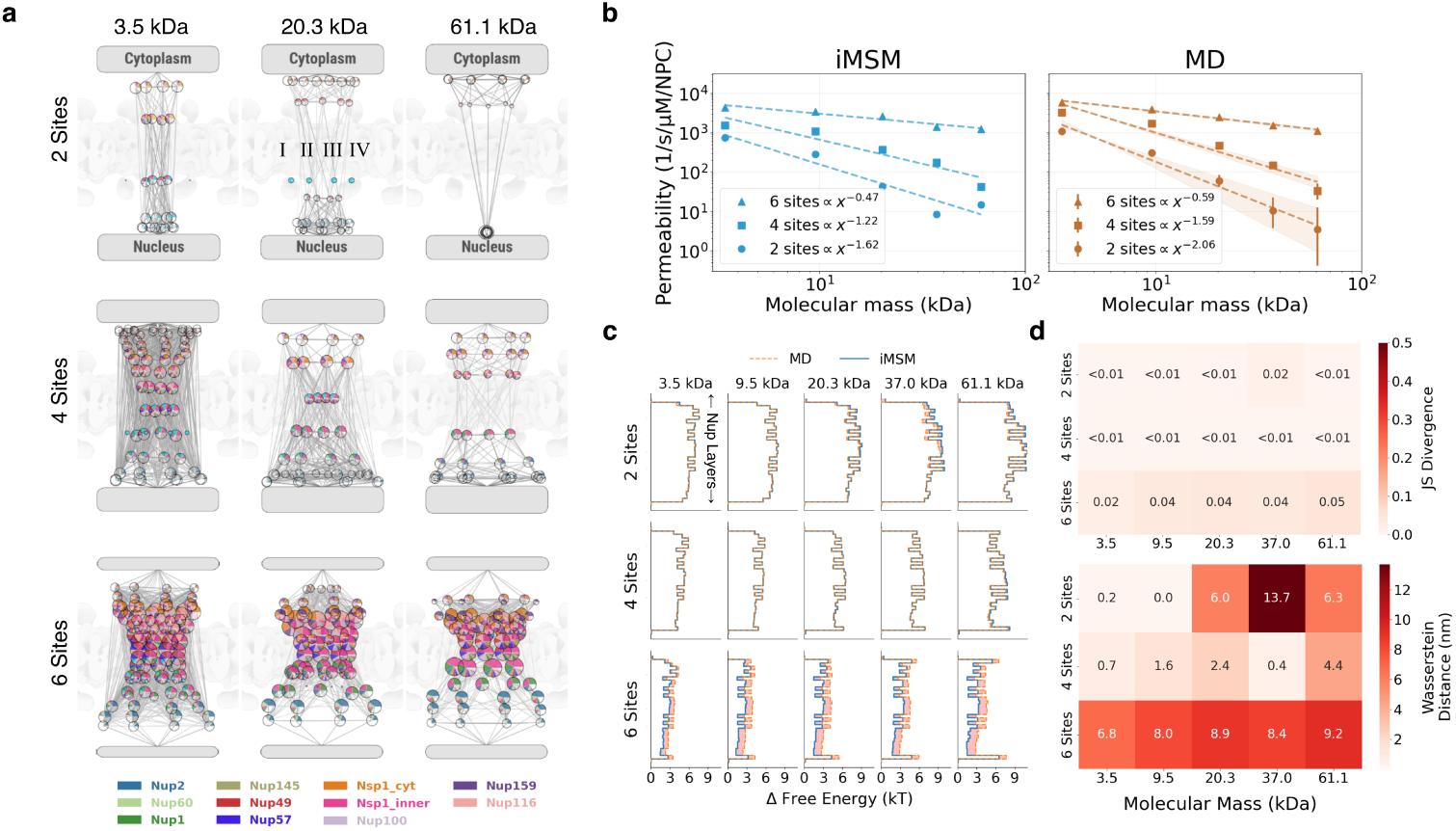
iMSMs reproduce transport properties across NTR size and interaction valency. **(A)** iMSMs for representative NTRs spanning three molecular masses and two, four or six FG-binding sites in a 54-nm-diameter NPC. Nodes represent interaction states, sectors indicate FG Nup composition, node size denotes stationary probability and edges represent state transitions. Cytoplasmic and nuclear unbound states are shown at the top and bottom, respectively. **(B)** Permeability estimated from iMSMs (left) and directly from MD simulations (right) for 15 NTR variants. Dashed lines show power-law fits for each valency, with fitted exponents indicated. Estimates used an aggregate MD simulation time of 1,800 *µ*s per variant. MD error bars, 95% Poisson confidence intervals. Shaded regions indicate the fit error. **(C)** Relative free-energy profiles across FG Nup layers, calculated from MD trajectories (orange, dashed) or iMSM stationary distributions (blue), for each molecular mass and valency. **(D)** JS divergence (top) and Wasserstein distance (bottom) between the corresponding MD and iMSM distributions. Wasserstein distances were calculated using the axial positions of the FG Nup anchor layers. Values are shown within each cell.

We first tested whether the accuracy observed for the representative NTRs extended across all 15 cargo variants at a pore diameter of 54 nm. iMSM-derived permeabilities reproduced the decrease in transport with increasing cargo size and the increase associated with higher FG-binding valency (Fig. 3b). Although absolute estimates differed from direct MD values by up to approximately half an order of magnitude, the relative trends were preserved across the full screen and at the other pore diameters examined, except for very sparsely sampled variants (Supp. Fig. S6).

The stationary distributions also remained close to their empirical MD counterparts, as indicated by consistently low Jensen–Shannon divergences and Wasserstein distances across all variants (Fig. 3c,d). Chapman-Kolmogorov tests across representative NTR variants and states likewise showed close agreement between predicted and observed multi-step transition probabilities, supporting the Markovian consistency of the iMSMs (Supp. Fig. S7). Thus, with sufficient input sampling, iMSMs preserve the principal kinetic and thermodynamic trends across a broad range of cargo sizes and interaction valencies.

### 2.4 iMSMs accelerate convergence of permeability estimates

We next tested whether iMSMs could estimate permeability using less input simulation time than direct transport tallying from MD. For each NTR variant, we trained iMSMs on progressively larger portions of the available trajectories and compared their permeability estimates with direct MD estimates obtained from the same data, using the full-dataset MD tally at *t* = 1800 *µ*s as the ground truth reference (Fig. 4a,b). The presented analysis uses progressively truncated trajectories, representing the practical setting of many short parallel simulations; results obtained by varying the number of full-length trajectories are shown in Supplementary Fig. S8. Full details of the subsetting procedures are provided in the Supplementary Methods.

**Fig. 4.**
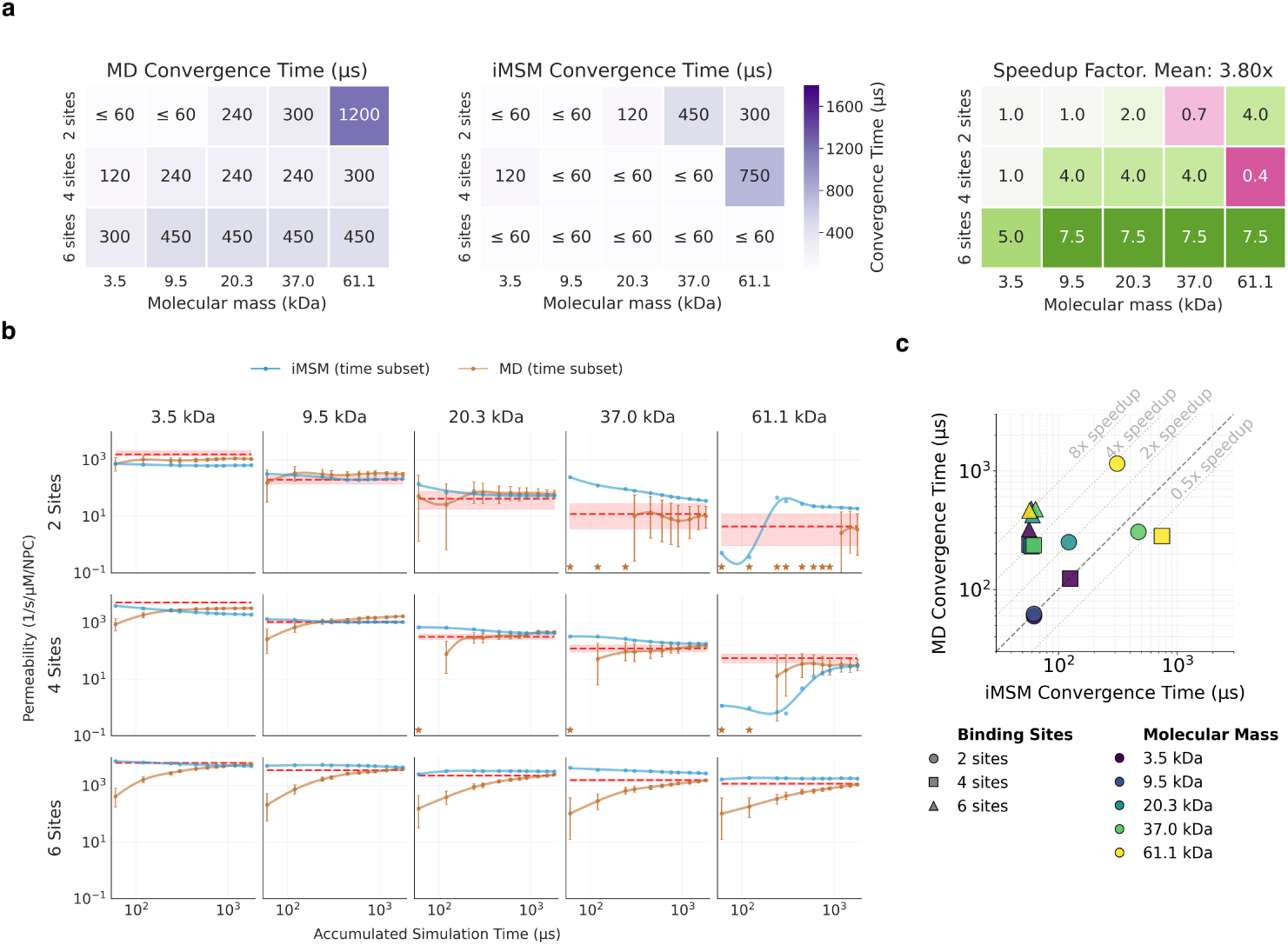
iMSMs accelerate convergence of permeability estimates. **(A)** Convergence times obtained by direct MD transport tallying (left) and iMSMs (center) for NTRs spanning five molecular masses and two, four or six FG-binding sites. Values indicate the earliest accumulated simulation time at which the permeability estimate entered and subsequently remained within twofold of its reference value. Corresponding MD-to-iMSM speedup factors are shown at right; the mean speedup across all variants was 3.80-fold. Values of *≤* 60 *µs* indicate convergence at or before the shortest accumulated simulation time evaluated. **(B)** Permeability estimates from iMSMs (blue) and direct MD tallying (orange) as a function of accumulated simulation time. Red dashed lines show permeabilities obtained from full time MD power-law fits across molecular masses, fitted separately for each number of FG-binding sites; shaded regions indicate the fit error. Orange error bars show 95% Poisson confidence intervals. Stars indicate subsets with zero observed transport events. **(C)** MD convergence time plotted against iMSM convergence time. Marker shape denotes the number of FG-binding sites and color denotes molecular mass.

For most NTR variants, iMSM-derived permeability converged to the full-data reference using less input simulation time than direct MD tallying (Fig. 4a,b). Using convergence to within a two-fold range of the reference permeability, iMSMs achieved an average speedup of 3.80 and a median speedup of 4.00 across all NTR variants. The benefit increased with NTR valency: average speedups for NTRs with 2, 4 and 6 interaction sites were 1.74×, 2.68× and 7.00×, respectively. Thus, iMSMs provided the largest acceleration for highly multivalent NTRs in the truncated-trajectory analysis. The mean speedup was smaller, 1.13×, when varying the number of full-length trajectories (Supplementary Fig. S8). The smaller gains for lower-valency variants likely reflect sparser sampling of informative interaction states, suggesting that targeted sampling within the central channel could further improve convergence.

### 2.5 Graded interaction states across pore diameters

We next used pore dilation as a controlled perturbation of the interaction-state networks and transport selectivity.

Previous studies have established that NPC diameter is not fixed. Structural and live-cell measurements have documented dilation and constriction associated with transport state and with mechanical or physiological perturbations, although the specific contribution of pore geometry to transport can be difficult to isolate in cells ([20], [67], [68], [69]). Experiments and simulations with FG-lined artificial nanopores show that wider pores accelerate Kap95 and BSA translocation and can substantially alter transport selectivity ([70], [71]).

Our previous study also examined the dependence of transport on pore diameter and cargo molecular weight using a simplified cylindrical pore containing 32 Nsp1 chains and comparing passive with facilitated diffusion [7]. The full NPC model allows us to examine the joint effects of pore diameter, cargo molecular weight, and discrete

NTR FG-binding valency on the interaction-state networks. Here, we vary these three parameters in a factorial design to determine how they individually and jointly shape transport thermodynamics and kinetics.

#### Pore dilation weakens size filtering in a valency-dependent manner

First, we calculated the free energy barrier magnitude (Δ*G*) at the pore center, *z* = 0, for all combinations of these parameters. Δ*G* increased with molecular weight across all pore diameters, indicating a size-dependent penalty for occupying the middle of the channel (Fig. 5a). However, the increase was steepest in narrow pores and became progressively weaker as the pore diameter increased, as demonstrated in the slope-versus-pore-diameter analysis. Thus, wider pores reduce the molecular-weight dependence of the central energetic barrier.

**Fig. 5.**
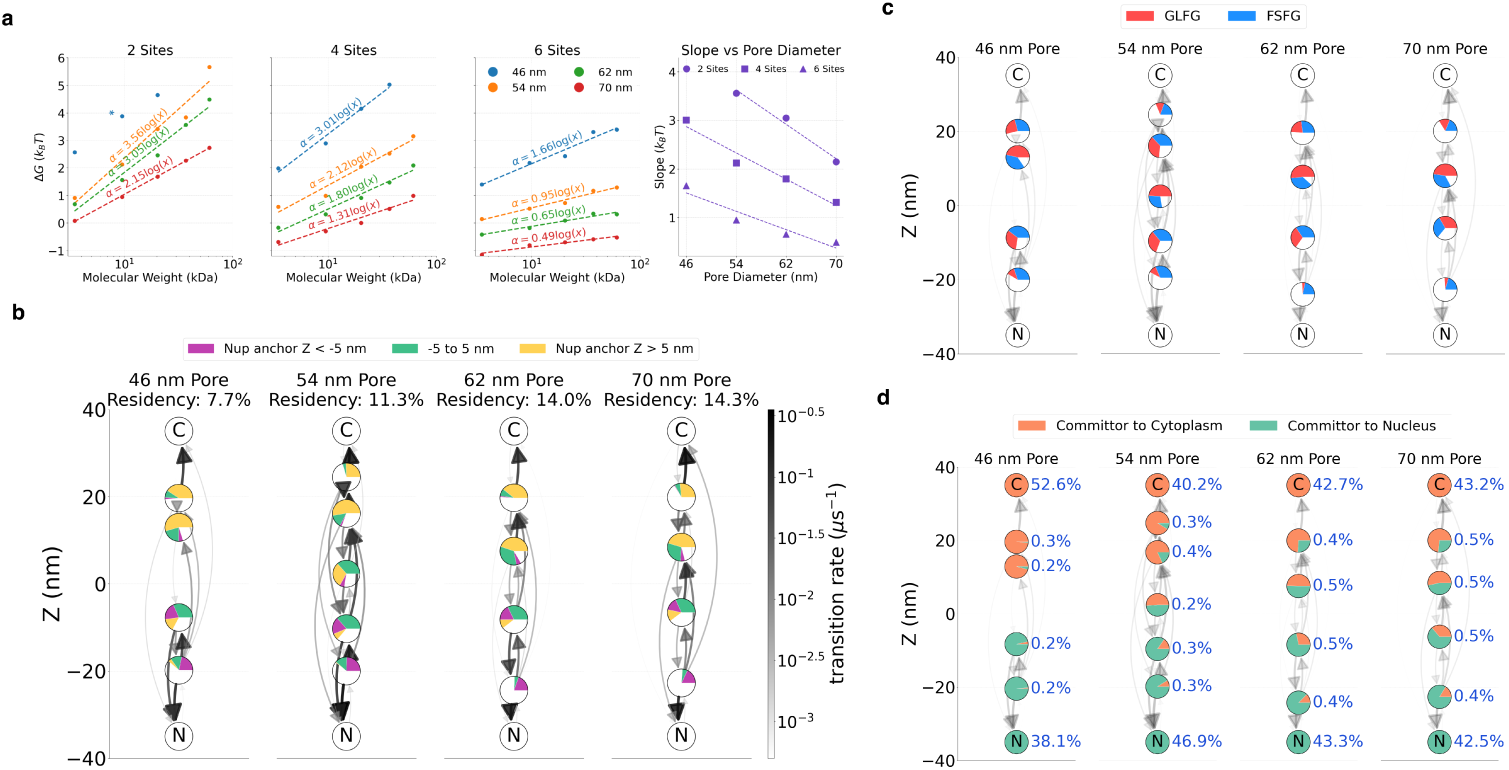
Pore dilation weakens size filtering while retaining broadly similar FG-mediated interaction environments. **(A)** Free-energy difference at the pore center, Δ*G*(*z* = 0), as a function of NTR molecular weight for receptors with two, four or six FG-binding sites in pores of the indicated diameters. Dashed lines show logarithmic fits, Δ*G* = *α* log(MW), with fitted slopes, *α*, indicated. Right, *α* as a function of pore diameter for each NTR valency. **(B–D)** Single-spoke iMSMs for a 20.3-kDa NTR with four FG-binding sites across the indicated pore diameters. Nodes represent interaction states and are positioned according to their estimated axial coordinate, *z*. Arrows represent state transitions, with arrow width and opacity indicating transition rate. C and N denote the cytoplasmic and nuclear unbound states, respectively. **(B)** Node wedges are colored according to the axial positions of the anchor sites of their constituent FG Nups. White wedges represent unoccupied reference interaction capacity. Residency indicates the combined stationary probability of the displayed interacting states. **(C)** Node sectors indicate the relative contributions of GLFG- and FSFG-containing Nups. **(D)** Orange and green sectors indicate committor probabilities towards the cytoplasmic and nuclear basins, respectively. Blue labels indicate state stationary probabilities.

This effect was further modulated by NTR valency. Low-valency NTRs showed the strongest molecular-weight dependence, particularly in narrow pores, whereas higher-valency NTRs exhibited shallower slopes (Fig. 5a). Pore dilation therefore weakens size-dependent exclusion, while multivalent FG-Nup engagement partially compensates for the steric or entropic cost of occupying the pore center. Both findings are consistent with our previous results from simplified pores in which permeability decreased more sharply with molecular weight for narrower pores and for more passive molecules [7].

#### Dilation modulates connectivity among graded interaction states

To determine how dilation changes the underlying transport mechanism, we constructed independent iMSMs for 20-kDa, four-site NTRs across pore diameters. These models retained the previously established agreement with MD permeability estimates (Supp. Fig. S6), and identified broadly similar interaction-state compositions across pore widths (Fig. 5b).

These independently constructed networks suggest that dilation primarily changes transition rates and connectivity among broadly similar interaction environments.

Similarly, for all pores, Nups anchored near the inner ring, particularly inner Nsp1, extended into cytoplasmic- and nuclear-side states (Supp. Fig. S9).

Wider pores also contained more open or weakly interacting states (Fig. 5b), consistent with lower time-averaged FG-Nup engagement. Partially engaged states combine FG-contact and unoccupied-capacity contributions within a finite time window. Their faster transitions to fully disengaged states, compared with strongly engaged states, identify kinetic routes from persistent multivalent binding towards release. State compositions and transition connectivity together describe these routes through graded engagement rather than reducing transport to a single bound-to-unbound step.

The spatial pattern of GLFG and FSFG contributions (as previously described for the representative NTR variants) remained broadly conserved across pore diameters (Fig. 5c). In particular, FSFG motifs remained more prominent on the nucleoplasmic side, and no systematic redistribution between motif classes was observed with increasing pore diameter. Thus, dilation primarily alters connectivity among states without redirecting transport toward a different FG-motif class.

Finally, we used committor analysis to examine the kinetic bottleneck for complete translocation. Committor values transitioned from cytoplasm-committed states on the cytoplasmic side to nucleus-committed states on the nucleoplasmic side (Fig. 5d). This transition became less sharp with increasing pore diameter, indicating that dilation smooths the kinetic barrier between cytoplasmic and nucleoplasmic commitment.

Overall, we found that pore dilation weakens the size-dependent central barrier while retaining broadly similar FG-mediated interaction environments. Within this model, dilation modulates selectivity through changes in confinement, state connectivity, and kinetic commitment within a broadly similar interaction-state organization.

### 2.6 iMSMs of the Slide-and-exchange mechanism

We next applied iMSMs to a molecular dynamics trajectory of Kap95 interacting with an Nsp1-FSFG125 construct containing six FSFG motifs, previously analyzed in [7] (Fig. 1c). To describe the motion of an individual FG motif across the receptor surface, we defined the Kap95 backbone sites as the interacting components and the mean C*α* position of a selected FSFG motif as the focal entity (Fig. 6a, Supp. movie 2).

**Fig. 6.**
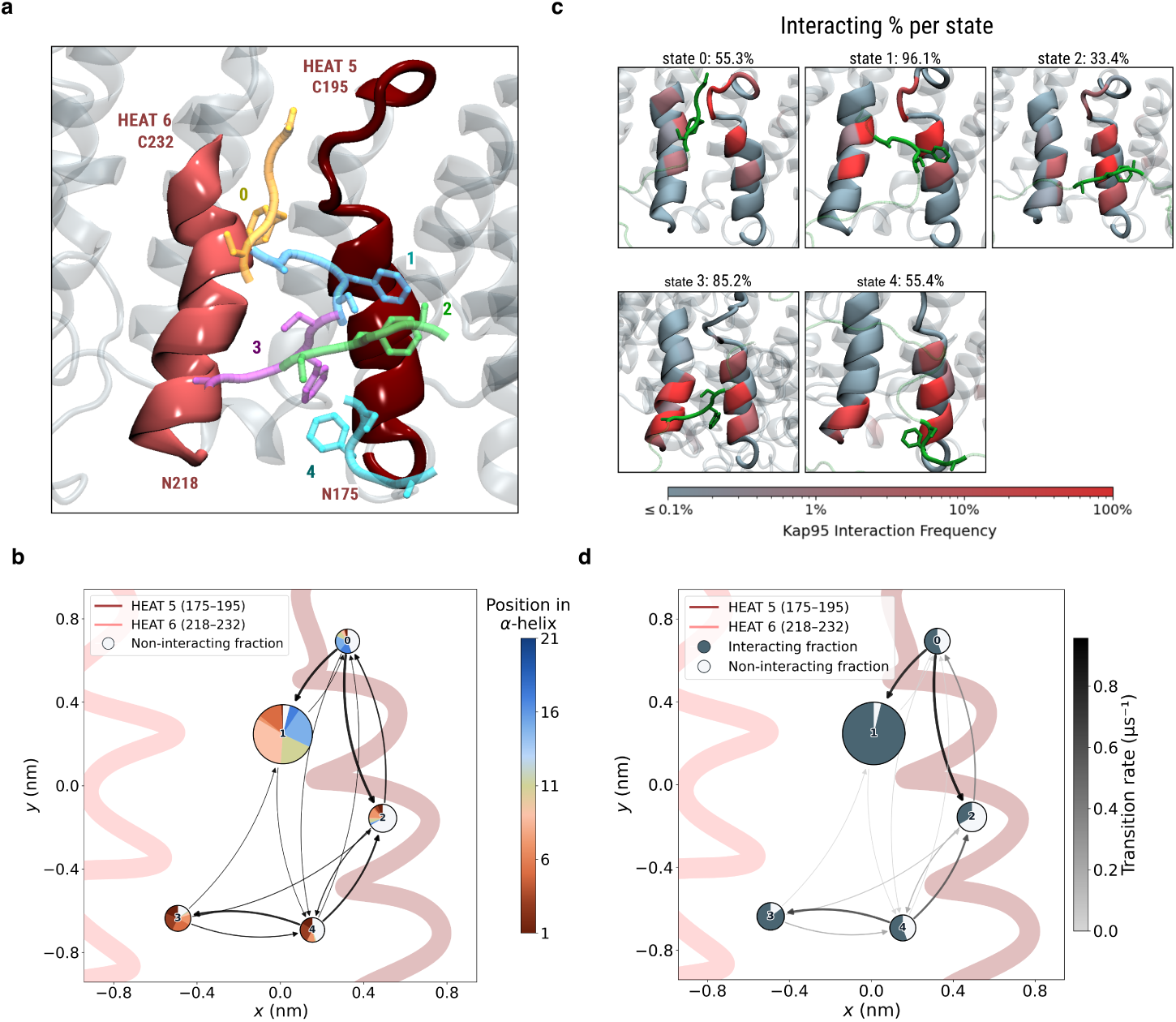
(A) Structural representation of the five iMSM states (0–4) adopted by an FSFG motif within the interhelical groove between HEAT repeats 5 (dark red) and 6 (light red) of Kap95. Representative FSFG configurations are colored by state and ordered according to the *y* coordinate of the motif center of mass. The remainder of the Nsp1-FSFG125 chain is omitted for clarity and other regions of Kap95 are shown in gray. Representative configurations were selected based on proximity to the cluster centers. **(B)** Spatially projected iMSM of FSFG motion along the HEAT 5/6 groove. Nodes represent recurring interaction states and are positioned according to the weighted mean *x* and *y* coordinates of the interacting residue backbones. Colored node wedges show the relative contributions of interacting Kap95 residues, colored by their mean *y* coordinate; white wedges show the unoccupied fraction of the reference interaction capacity. Arrows indicate observed transitions between states. Node size is proportional to stationary probability. **(C)** Kap95 contact-frequency maps for states 0–4. Kap95 residues are colored according to their interaction frequency with the FSFG motif within each state, using the logarithmic scale shown; the FSFG motif is shown in green. Percentages above the panels report the mean fraction of the reference interaction capacity occupied by Kap95 contacts in each state. **(D)** Kinetic representation of the same iMSM. Filled and white node wedges show the occupied and unoccupied fractions of the reference interaction capacity, respectively, and arrow shading indicates transition rates according to the scale shown. Together, the states and their connectivity show that the FSFG motif moves anisotropically along the Kap95 groove by exchanging residue-level contacts while remaining continuously associated with the receptor. This iMSM defines each state by the time-averaged distribution of contacts between one focal FSFG motif and Kap95 residues, thereby grouping many microscopic geometries that share the same local interaction pattern; consequently, state transitions report reorganization of the motif’s local interaction environment rather than changes in the global protein conformation.

The resulting iMSM resolves recurrent interaction states distributed along the inter-helical groove formed by the HEAT repeats of Kap95 (Fig. 6b). Transitions between these states resolve anisotropic motion of the FG motif along the groove (Fig. 6b,d), consistent with previously proposed slide-and-exchange dynamics [5, 72, 73].

Notably, the FSFG motif remained associated with the HEAT 5/6 groove throughout the simulation, while the fraction of the reference interaction capacity occupied by Kap95 contacts varied substantially among iMSM states (Fig. 6d). Thus, movement along the groove does not require complete dissociation. Instead, the motif moves between different degrees of engagement, exchanging individual residue-level contacts and moving between neighboring binding configurations, a slide-and-exchange mechanism.

## 3 Discussion

Across the two molecular resolutions studied here, iMSMs reveal a common organization of transport kinetics into graded interaction states. At the receptor surface, FG motifs exchange residue-level contacts while remaining associated with Kap95. At the pore scale, partially engaged states provide faster routes toward disengagement than strongly bound states. Together, these results provide a quantitative kinetic picture of how dynamic multivalent systems can sustain strong interactions while retaining kinetic routes out of strongly bound states.

The method is demonstrated here for nucleocytoplasmic transport across two distinct scales. However, the same representation may be useful wherever kinetics can be described through recurring local interaction environments around user-defined focal entities. This abstraction extends naturally to condensate biology, where multivalent interactions, compositional heterogeneity, and exchange kinetics are central features [74, 75]. Stress granules, for example, contain stable and dynamic substructures whose multistep assembly and disassembly motivate a local kinetic description [74, 76–79]. Focal entities could follow scaffold or client molecules, with states describing local engagement, composition and exchange between recurring environments, consistent with the structural and kinetic heterogeneity of stress granules [76, 77]. For LAT signaling condensates, focal entities could follow LAT molecules or microclusters to describe local interaction dynamics [9, 80]. In transcription-associated condensates, focal entities could follow nascent RNA or local Pol II/cofactor environments, where RNA can promote and later dissolve condensates [81, 82].

These prospective applications suggest a useful boundary for the method: iMSMs are most compelling when the process of interest can be represented in terms of recurrent local interaction states and their kinetic coupling, and when the resulting state decomposition is sufficiently Markovian at an appropriate lag time [43]. In practice, this should be assessed using standard MSM validation criteria, such as implied-timescale convergence [62] and Chapman-Kolmogorov consistency tests [43]. Mesoscopic observables such as droplet number or mean droplet size may still be incorporated, but most naturally as part of a broader state description rather than as the sole variable of the model. To model more complex behaviors, we suggest an extension to the iMSM framework, which is based on the idea of multiple focal points within a single system (supplementary information).

State identification and transition estimation are automated, but the physical representation and temporal resolution determine what the model can resolve. The resulting networks support mechanistic interpretation of the simulated dynamics rather than establishing a unique causal explanation. Validation across different NTR variants tests robustness, but does not quantify uncertainty for individual systems. This could be computed, for example, through Bayesian iMSMs with a Dirichlet prior [83], yielding posterior distributions over transition probabilities, or through a bootstrap procedure comprising random sampling with replacement from the pool of MD trajectories.

Once validated, iMSMs provide compact kinetic fingerprints that enable systematic, high-throughput comparison across large simulation datasets spanning many perturbations. Differences between these fingerprints can reveal how each perturbation reshapes interaction states, pathways, and kinetics. For example, an FG-Nup deletion can be evaluated by constructing a comparable iMSM and quantifying its divergence from the wild-type transition matrix. This suggests an engineering pipeline: define the wild-type kinetic target, evaluate candidate NTR designs in the perturbed environment, and optimize them to reduce kinetic divergence while also monitoring macroscopic properties such as permeability. Unlike a single scalar observable, the transition matrix retains state- and pathway-specific kinetic information. Realizing this approach will require compatible state representations, uncertainty-aware comparison metrics, and independent validation that restored kinetics translate into restored function, but could ultimately turn mechanistic models of complex biomolecular dynamics into quantitative objectives for biomolecular design.

The probabilistic structure of iMSMs, complemented by uncertainty estimates, would also make them natural inputs to Bayesian metamodeling, which couples heterogeneous models through shared probabilistic representations [84, 85]. For example, an iMSM of transport through a single nuclear pore could be coupled to a whole-cell transport model [20] to predict transport under non-equilibrium conditions, following the multiscale strategy used previously [7]. More broadly, coupling iMSMs of different cellular processes could provide building blocks for whole-cell models that connect local molecular interactions to the interplay between processes and, ultimately, to the behavior of the cell as a whole.

## Funding

This work was supported by the Israel Science Foundation (grant no. 385/24) and by a Minerva Center Grant on Cell Intelligence.

## Code availability

The repository containing the generic implementation of the interaction-based Markov state model (iMSM) framework, the NPC-specific implementation used in this study, and the code for reproducing the figures is available at https://github.com/ravehlab/iMSM.

## S1 Supplementary Information

### S1.1 Supplementary Methods

#### S1.1.1 Generating a molecular dynamics dataset of nucleocytoplasmic transport

Nucleocytoplasmic transport trajectories were generated using the coarse-grained Brownian dynamics model of the yeast nuclear pore complex (NPC) described previously [7]. Briefly, the model represents the experimentally informed NPC scaffold together with 216 intrinsically disordered FG-Nup chains belonging to ten FG-Nup types. The FG-Nup domains were anchored at their corresponding positions in the NPC scaffold and modeled using one bead per 20 amino acids. Interactions between FG-Nup beads, transported particles and the NPC scaffold were described using custom force fields, with parameters and implementation as reported previously [7]. Unless stated otherwise, all model parameters were unchanged from that study.

Transported nuclear transport receptors (NTRs) were represented as spherical particles with molecular masses of 3.5, 9.5, 20.3, 37.0 or 61.1 kDa. Each NTR carried 2, 4 or 6 FG-binding sites, producing 15 combinations of molecular mass and interaction valency. Binding sites were distributed uniformly over the NTR surface. For each NTR variant, simulations were performed in NPCs with central-channel diameters of 46, 54, 62 or 70 nm. Pore dilation was modeled by radially scaling the NPC ring coordinates while preserving the scaffold stoichiometry and the total number of FG-Nup chains. Consequently, FG-Nup density decreased as pore diameter increased.

For each combination of FG-binding sites and pore diameter, 30 independent simulations were initialized with 100 copies of each NTR mass. Initial configurations were subjected to an equilibration period of 10 *µs*.

Each production trajectory had a duration of 70 *µ*s for 54 nm diameter NPCs, and 30 *µ*s for the rest. After excluding the first 10 *µ*s as equilibration, the analyzed production durations were 60 and 20 *µ*s, respectively. Coordinates and interaction data were recorded every 100 ns. Across 30 independent trajectories, this yielded an aggregate production time of 1.8 *m*s for each NTR–pore condition for 54-nm-diameter NPCs and 0.6 *m*s for the rest. In total, the complete dataset comprised 12 simulation variants, 360 independent trajectories and 10.8 *m*s of Brownian dynamics.

#### S1.1.2 General construction of iMSMs

An iMSM requires three modeling choices: the interacting components to be resolved, the focal entity whose environment is followed, and the features used to describe that environment. The system particles are first partitioned into a disjoint set of components, *C* = {*c*_1_*, … , c_N_*}. Depending on the spatial and mechanistic resolution required, a component may represent an atom, interaction site, protein domain, complete protein, molecular complex or a defined group of these elements. A focal entity *f* is then selected. It may coincide with an individual component, be derived from a group of components, or be fixed at a spatial location. The resulting model describes transitions between recurring environments around *f*, rather than transitions between global conformations of the full system.

For each input MD trajectory, we compute an interaction trajectory *I*(*t*) describing the relationship between the focal entity and the components in *C*. In the present NPC application, the primary features are spatial-proximity interactions between individual transported NTRs and neighboring FG-Nup components. The framework is not restricted to binary contacts: interaction features may be supplemented by consistently measurable local or coarse-grained observables. Examples include interaction-site saturation, the composition or number of nearby component types, local density, the fraction of a component in a dense phase, condensate size, or other regional observables relevant to the process under study.

To reduce sensitivity to stochastic contact formation and rupture, the interaction trajectory *I*(*t*) is segmented into consecutive windows of duration *τ*. For each focal entity *f*, we define a per-frame interaction capacity *C_f_*, which specifies the total interaction weight that can be represented at each sampled time point. Let *n_i_*(*t*) denote the interaction contribution of component *c_i_* to the focal entity at time *t*. The remaining unoccupied interaction capacity at that time point is

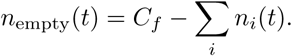

For a time window *W* containing *N_W_* sampled frames, the interaction histogram *H* is constructed by accumulating the interaction contributions over all frames and normalizing them by the total interaction capacity available over the window. Thus, the contribution of component *c_i_* is

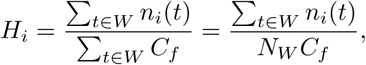

and the unoccupied contribution is

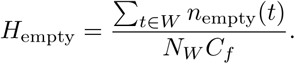

Consequently, the complete interaction histogram sums to one while retaining both the identities of the interacting components and the overall degree of engagement of the focal entity during the window.

Importantly, this normalization differs from normalizing by the total number of interactions actually observed within the window, ∑*_t_*_∈W_ ∑*_i_* n*_i_*(*t*). Such a normalization would retain only the relative composition of the observed interactions and would discard differences in their overall magnitude. For example, two windows involving the same relative proportions of interaction partners would appear identical even if the focal entity were strongly engaged throughout one window but only weakly engaged during the other. Normalization by the total available interaction capacity instead preserves this distinction, allowing iMSM states to resolve strongly, partially, and weakly engaged interaction environments.

Additional consistently measurable observables may be appended to the interaction histogram when required to resolve process-specific dynamics. The resulting window-level feature vectors are clustered to produce the reduced state space *S* = {*s*_1_*, … , s_k_*} of recurring local interaction environments.

Kinetic evolution is estimated by counting transitions between the resulting state assignments at lag time *τ* and normalizing these counts to obtain the transition-probability matrix *T*. Each element *T_ij_* gives the probability of observing state *s_j_* after one lag time when the current state is *s_i_*. Standard MSM observables can subsequently be computed from *T*, including stationary probabilities and trajectory-derived kinetic quantities.

The same construction can be applied beyond cargo-centered NPC transport when a process can be described through recurring local interaction environments. For example, in a phase-separating system, focal entities could be attached to representative molecules or fixed spatial points, allowing states to encode local interaction composition together with observables that distinguish dilute, dense or interfacial environments. Such applications require system-specific choices of components, features and lag time, followed by the same state-discretization and transition-estimation procedure.

An open-source implementation of the iMSM framework is available at https://github.com/xroi/iMSM.

#### S1.1.3 Construction of iMSMs of Nucleocytoplasmic Transport

The NPC model contained 216 individual FG-Nup chains belonging to ten distinct FG-Nup types. To represent differences between the relatively constrained region near each anchoring site and the more mobile region of the disordered domain, each FG-Nup chain was divided into two components corresponding to its N- and C-terminal halves. Each NTR variant, differing in molecular size, interaction valency or both, was defined as a separate component. Individual transported NTRs also served as the focal entities whose interaction environments were followed when constructing the iMSMs.

For each focal NTR, we constructed an instantaneous interaction vector describing its contacts with the FG-Nup components. All NTR variants were represented using the same per-frame reference interaction capacity, *C*_NTR_ = 5. This quantity defines the normalization of the iMSM interaction representation and is distinct from the two, four, or six FG-binding sites specified for the NTRs in the underlying simulation model.

At each sampled time point, the interaction contributions of the FG-Nup components occupied part of this capacity, while any remaining capacity was represented explicitly as unoccupied. To retain information about the spatial location of an incompletely engaged NTR, the unoccupied contribution was assigned to one of five spatial components according to the instantaneous NTR *z*-coordinate: cytoplasm (*z* ≥ 15 nm), cytoplasmic channel (5 *< z <* 15 nm), inner channel ( 5 *z* 5 nm), nuclear channel ( −15 *< z <* − 5 nm), or nucleus (*z* 15 nm). These spatially resolved unoccupied quantities are components of the interaction representation rather than predefined iMSM states; the final states are obtained subsequently by clustering the window-level feature vectors.

The interaction trajectories were partitioned into consecutive 5 *µ*s windows. For each window, interaction contributions were summed over all sampled frames and normalized by the sum of the available interaction capacities over those frames. Because *C*_NTR_ = 5 at every frame, a window containing *N_W_* sampled frames has a total interaction capacity of 5*N_W_*. The contribution of FG-Nup component *i* to the window-level histogram was therefore calculated as

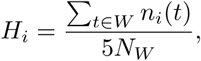

with the remaining fraction corresponding to the spatially resolved unoccupied capacity. This normalization ensures that each interaction histogram sums to one while retaining differences in the total degree of FG-Nup engagement. In particular, windows with similar relative FG-Nup composition remain distinguishable when they differ in the fraction of the available interaction capacity that is occupied. The mean NTR *z*-coordinate within each window was appended to the interaction histogram as an additional clustering feature. To control its contribution relative to the interaction components, the mean *z*-coordinate was multiplied by a fixed weight,

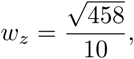

where 458 was the feature count used to set the relative scale in the implementation. The same weighting was applied to every NTR and pore condition.

Window-level feature vectors were clustered using *k*-means with an initial resolution of *k* = 320. Assignment to the nearest cluster center used exact squared-Euclidean distance, implemented with the FAISS IndexFlatL2 index. Because squared Euclidean distance preserves the ordering of Euclidean distances, this is equivalent to assigning each feature vector to its nearest center under the Euclidean metric.

The value *k* = 320 was selected empirically after examining models constructed at several clustering resolutions. Lower resolutions increasingly combined interaction environments that were useful to distinguish mechanistically, whereas higher resolutions produced progressively sparser states without appreciably improving the recovery of the principal stationary and kinetic observables. We therefore selected 320 as a common operating resolution that balanced state interpretability, sampling, and model accuracy. This value was fixed before the comparative analyses and was not tuned separately for different NTR variants or pore diameters.

We subsequently consolidated redundant or weakly sampled clusters. First, clusters whose centers assigned more than 0.95 of their interaction-histogram weight to the nuclear unoccupied component were merged into a single canonical nuclear state; clusters satisfying the analogous criterion for the cytoplasmic unoccupied component were merged into a single canonical cytoplasmic state. Second, near-duplicate states were merged when the *L*_1_ distance between their feature-space cluster centers was less than 0.005:

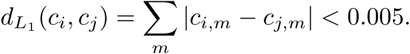

Finally, states represented by fewer than ten window assignments in the input trajectories were excluded from the final model. Consequently, 320 denotes the common initial clustering resolution, while the number of retained iMSM states could be smaller and could differ among NTR and pore conditions.

The transition models used a lag time of 5 *µ*s, corresponding to 50 trajectory-sampling intervals of 100 ns. This lag time was selected on the basis of implied-timescale convergence analysis (Supplementary Fig. S1).

A pseudocount of 10^−3^ was added to each element of the transition matrix before row normalization.

#### S1.1.4 Incorporation of NPC rotational symmetry

The NPC scaffold has eightfold rotational symmetry, such that configurations separated by rotations of 45^◦^ around the transport axis are physically equivalent. We incorporated this symmetry at both the trajectory-augmentation and clustering stages. Each transport trajectory was transformed into eight symmetry-equivalent copies by applying rotations of 0^◦^, 45^◦^*, … ,* 315^◦^ around the central axis of the NPC.

We then adapted the *k*-means clustering procedure so that interaction states were identified as sets of eight symmetry-related clusters (Supp. Fig. S2). For every candidate cluster center, the seven corresponding centers generated by successive 45^◦^ rotations were included in the state representation. Assignment and center estimation were therefore constrained such that symmetry-equivalent interaction environments were treated consistently across the eight NPC spokes. This procedure increased the effective amount of sampling available for state estimation without treating rotationally equivalent transport events as distinct physical mechanisms.

#### S1.1.5 Lag-Time Selection

We selected the iMSM lag time by examining the implied timescales obtained over a range of candidate lag times (Supp. Fig. S1). The dominant implied timescales approached a plateau at approximately 50 sampling steps, corresponding to 5 *µ*s given the trajectory-sampling interval of 100 ns. We therefore used 5 *µ*s as the lag time for the analyses presented in this study.

This analysis differs from a conventional implied-timescale test for an MSM with a fixed state discretization. In the iMSM framework used here, the identified interaction states depend on the selected lag time as a histogram is calculated over this period to characterize the state. Consequently, for each candidate lag time, the interaction states were regenerated and the corresponding transition matrix was estimated independently. The curves in Supp. Fig. S1 therefore do not necessarily represent identical dynamical modes evaluated over an unchanged state space. Rather, the analysis tests whether the dominant kinetic timescales inferred by the complete lag-dependent iMSM construction become broadly stable as the lag time increases. Their approximate stabilization near 50 sampling steps supported this choice as a balance between kinetic consistency and retention of sufficient transition statistics.

#### S1.1.6 State Position Estimation

For the purposes of visualization (Figs. 1 and 2), we estimated the three-dimensional position of each iMSM state through a heuristic based on the anchor positions of the Nups composing the states. First, we assigned each iMSM component (i.e., a specific Nup) to a three-dimensional volume in the simulation based on its anchor position.

Let *u_i_* denote the center of mass of the volume for component *i*, and let *p_i,j_* represent the probability of component *i* in the histogram of state *j*. The estimated 3D position of state *j*, denoted as *U_j_*, is calculated as the mean of these centers of mass, weighted by their respective probabilities:

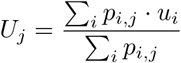

#### S1.1.7 Permeability Estimation

The permeability of NTRs in the simulation was estimated in two ways: through explicit transport tallying directly from the MD trajectories, and by simulating synthetic trajectories within the iMSM state space to tally transport events.

##### Explicit transport tallying in MD

We defined two spatial regions: the free cytoplasm and NPC-bound states on the cytoplasmic side (*z ≥* 10 nm), and the free nucleus and NPC-bound states on the nucleoplasmic side (*z ≤-*10 nm). We explicitly tallied the number of successful transport exchanges between these regions directly from the MD trajectories. The final permeability value was calculated as the total number of transport events divided by the total simulated time, the NTR concentration, and the number of NPCs (which is 1 in our simulation):

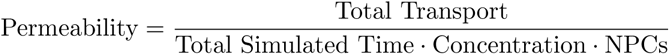

Confidence intervals for these estimates were calculated assuming the transport events follow a Poisson distribution.

##### Transport tallying in iMSMs

As an orthogonal approach, we estimated permeability by generating trajectories in state space. We initially defined two groups of iMSM states based on the z feature of the state: *S_C_*, comprising cytoplasmic states (*z ≥*10 nm), and *S_N_*, comprising nucleoplasmic states (*z ≤-*10 nm). We simulated 10,000 trajectories from the iMSMs, each consisting of 1,000 time steps, with the core objective of tallying the successful transport passages between *S_C_* and *S_N_*.

To improve the robustness of our estimation, we modified the strict spatial boundaries used to define these states by introducing a 5 nm buffer region. Instead of relying on a binary cutoff, we assigned a continuous weight to each state based on its estimated z-axis position. Specifically, a state’s weight starts at 0 at |*z*| = 5 nm, grows linearly to 1 at |*z*| = 10 nm, and remains 1 for all regions where |*z*| ≥ 10 nm.

Consequently, each transport event in the tally was scaled by the product of corresponding state weights. The permeability was then computed using the same equation as the MD tallying, where the concentration is that of a single molecule, the number of NPCs is 1, and the total simulated time is defined as the variable time step multiplied by the total number of steps (1,000).

#### S1.1.8 GLFG/FSFG Analysis

To characterize the FG-motif composition of each iMSM interaction state, we grouped FG Nups according to their relative contributions of GLFG- and FSFG-type motifs, following the motif composition used in the underlying NPC simulation model [7]. Nup100, Nup116, Nup49, Nup57, and Nup145 were assigned to the GLFG group, whereas Nup159, Nup60, and Nup2 were assigned to the FSFG group. Nsp1 and Nup1 contain contributions from both motif classes and were therefore partitioned between the two groups according to their motif composition in the simulation model. Specifically, Nsp1 was assigned weights of 0.3273 to GLFG and 0.6727 to FSFG, while Nup1 was assigned with weights of 0.3475 to GLFG and 0.6525 to FSFG.

For each iMSM state, the GLFG and FSFG contributions were calculated from the corresponding FG-Nup composition of the state. Thus, the total GLFG contribution was calculated as

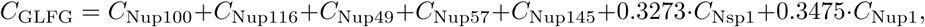

and the FSFG contribution as

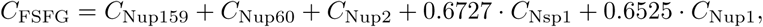

where *C*_Nup_ denotes the contribution of the indicated FG Nup to the interaction histogram of that state.

#### S1.1.9 Permeability Convergence Analysis

To evaluate the computational efficiency and acceleration provided by the iMSM framework compared to Molecular Dynamics simulations, we established the following metric to determine the convergence time of cargo permeability. For each NPC variant and cargo size, the full MD dataset comprised 30 independent simulations, each 60 *µ*s in length, yielding a maximum accumulated simulation time of 1800 *µ*s. Subsets of varying accumulated simulation times *t_k_* ∈ [60, 1800] *µ*s were constructed by truncating each of the 30 trajectories to a fraction of its total duration (ranging from 2*/*60 to 60*/*60). Permeability estimates for both the direct MD calculations and the iMSM predictions were then computed at each accumulated simulation time point.

iMSMs for short-duration subsets (2 *µ*s, 4 *µ*s, 8 *µ*s) used shorter time-averaged windows of 1 *µ*s, 2 *µ*s and 4 *µ*s respectively, instead of the standard 5 *µ*s for the longer duration subsets.

The convergence time, *t*_conv_, was defined as the earliest accumulated simulation time at which the estimated permeability enters and subsequently remains within a designated envelope around the final converged value:

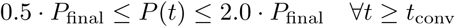

where *P*_final_ represents the permeability computed at the maximum accumulated simulation time of 1,800 *µ*s. Because the MD permeability is calculated directly from full transport counts divided by the total elapsed simulation time, it is inherently a cumulative average rate and was analyzed without further smoothing. In contrast, because the iMSM estimates represent discrete, independent model predictions at each trajectory length, they are subject to fluctuations between time points. To filter out this high-frequency noise and ensure a robust assessment of convergence, a running mean (cumulative average) of the iMSM estimated permeabilities was computed over the sorted simulation time subsets prior to applying the convergence criterion. The speedup factor was then computed as the ratio of the MD convergence time to the corresponding iMSM convergence time.

As a complementary subsetting strategy, we varied the number of independent full-length trajectories included in the analysis while retaining the complete 60 *µ*s duration of each selected trajectory. For a subset containing *n* trajectories, the accumulated simulation time was therefore 60*n µ*s. At each subset size, permeability was estimated both by direct MD tallying and by an iMSM constructed from the same selected trajectories. The resulting convergence behaviour is shown in Supplementary Fig. S8.

#### S1.1.10 Construction of the Kap95–FSFG iMSM

The Kap95–FSFG analysis used an all-atom molecular dynamics trajectory reported previously in [7]. The simulated system contained Kap95 together with two Nsp1-FSFG125 chains, each containing six FSFG motifs, and comprised approximately 14.2 *µ*s of production simulation. System preparation, force-field parameters, equilibration, and the Anton2 production-simulation protocol are described in detail in the previous study [7] and were not modified for the present analysis. Coordinates were sampled every 0.96 ns, and all available production frames were included without additional temporal subsampling or trajectory trimming.

For the present analysis, we selected the Nsp1-FSFG125 chain that remained associated with the inter-helical groove between HEAT repeats 5 and 6 of Kap95 during the trajectory; the second Nsp1 chain was not included in the iMSM construction. We focused on one FSFG motif within this associated chain. The focal position of the motif was represented by the arithmetic mean of the C*α* coordinates of its four constituent residues. Each of the 861 Kap95 C*α* sites was treated as a separate interacting component, such that the iMSM describes changes in the local Kap95 interaction environment sampled by the selected FSFG motif.

Interactions were defined using a distance threshold of 1.0 nm between the focal FSFG motif and Kap95 C*α* sites. Following the general iMSM construction described above, the focal entity was assigned an interaction capacity of five. At each sampled frame, up to the five Kap95 components interacting with the motif contributed to this capacity, while capacity not occupied by an interaction was represented as the non-interacting fraction. Thus, the representation retained both the identities of Kap95 sites contacted by the motif and the overall degree of engagement with the Kap95 surface.

The interaction trajectory was divided into consecutive windows of 500 simulation frames, corresponding to 480 ns per window. For each window, interaction contributions were accumulated over the constituent frames and normalized by the total available interaction capacity, 5×500. The resulting feature vector therefore describes the time-averaged distribution of contacts between the FSFG motif and individual Kap95 sites together with the fraction of unoccupied interaction capacity. This normalization allows windows involving similar regions of Kap95 to remain distinguishable when the motif differs in its overall degree of association.

Window-level interaction vectors were clustered using *k*-means with *k* = 5. Five states were chosen to provide an interpretable coarse-graining of the motif’s motion along the Kap95 binding groove while retaining distinct local interaction environments. No additional positional coordinate was included in the clustering; the states were defined solely from the interaction representation.

Transitions between state assignments separated by one 480-ns window were counted to construct the transition-count matrix. A pseudocount of 0.01 was added to each element of this matrix before row normalization to obtain the transition-probability matrix. The resulting five-state iMSM therefore describes the kinetics of exchange among recurring, time-averaged Kap95 interaction environments rather than transitions between individual instantaneous FSFG conformations.

### S1.2 Multiple focal entities extension

Capturing mesoscopic observables like droplet size inherently requires moving beyond the perspective of an isolated molecule. While the single-focal-entity iMSM framework successfully models these isolated local dynamics, such complex biomolecular phenomena are ultimately driven by the concerted action of multiple interacting centers. A natural extension of this approach involves simultaneously defining a set of *M* interaction focal entities, *F* = {*f*_1_*, … , f_M_}*. By tracking the evolving interaction environments across several focal entities at once, each exploring a local state space 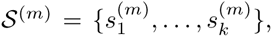, the iMSM can hypothetically be expanded to characterize not just independent local fluctuations, but collective dynamics spanning an entire macromolecular system. The global state of the system at time *t* is then represented by the full collection of local states, **S**(*t*) = (*s*^(1)^(*t*)*, … , s*^(*M*)^(*t*)). Furthermore, for systems containing equivalent molecules, or when the set of focal entities is defined on a uniform spatial grid, the local state spaces ^(*m*)^ and their underlying transition probabilities become invariant under compositional or translational symmetry. This allows interaction trajectories to be pooled across symmetric entities, potentially improving statistical sampling and state estimation.

Modeling multiple focal entities simultaneously requires extending the kinetic framework to capture the interdependencies between different local environments. In such an expanded model, the temporal evolution of **S**(*t*) can rely on two distinct mathematical operators: self-transition matrices 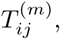, which describe the independent probability that focal entity *f_m_* transitions from state 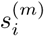 to state 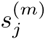 over lag time *τ*, and cross-transition matrices 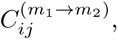 which quantify how the current state of focal entity 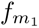 influences the subsequent state of 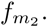 By combining these terms through a unifying coupling function *f*_couple_, the local transition probabilities take the form:

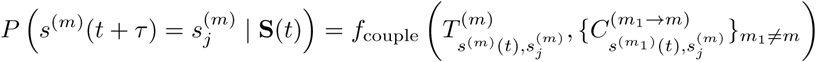

This multi-entity framework introduces some trade-offs. Calculating more interactions and transition matrices increases analysis time, though this overhead is likely negligible compared to the underlying MD simulations. Furthermore, relying on *f*_couple_ is a statistical estimate rather than a strictly rigorous global coupling. However, this approximation is a necessary and advantageous compromise; indeed, recent theoretical frameworks increasingly propose decomposing large systems into separable subsystems that can ultimately be coupled [48]. Constructing a true global state space is combinatorially explosive; by instead factorizing kinetics into localized TMs and CTMs, transition probabilities converge much faster, potentially rendering large-scale collective dynamics statistically tractable.

By explicitly coupling these spatial and temporal transition terms, an extended iMSM may model complex, distributed kinetic behaviors, such as the communication between stable and dynamic substructures in stress granules, the collective coordination of LAT microclusters during T cell signaling, or the RNA-driven feedback that restructures transcription-associated condensates.

### S1.3 Supplementary Movies

***Supplementary Movie 1. From molecular simulation to interaction-based Markov state models of nucleocytoplasmic transport.***

The movie begins with a visualization of the coarse-grained yeast NPC simulation system described previously [7], and transitions to the corresponding iMSM for a 61.1 kDa NTR with four FG-Nup interaction sites, as shown in Fig. 2a. Nodes represent recurring interaction states, with sectors indicating their FG-Nup composition, while edges indicate transitions between states. The iMSM is animated using Monte Carlo trajectories generated from its transition-probability matrix, illustrating stochastic transport between the cytoplasmic and nuclear unbound states through successive FGNup interaction environments. The movie then transitions to the collection of iMSMs shown in Fig. 3a, illustrating how the interaction-state networks vary across NTR molecular masses and FG-binding valencies.

***Supplementary Movie 2. Mapping FSFG–Kap95 interactions onto interaction-based Markov states.***

The movie shows a side-by-side representation of the molecular dynamics trajectory and its corresponding iMSM, as analyzed in Fig. 6. In the left panel, Kap95 is shown in gray and Nsp1-FSFG125 in green; Kap95 residues interacting with the selected FSFG motif in each simulation frame are highlighted in red. In the right panel, the corresponding iMSM is shown, with the current interaction state highlighted as the trajectory progresses, illustrating how the evolving residue-level interaction pattern is mapped onto discrete iMSM states. Node sectors indicate the interacting and non-interacting fractions associated with each state, and edges indicate transitions between states, with transition rates shown according to the accompanying scale. Together, the two views illustrate how continuous rearrangement of FSFG–Kap95 contacts in the molecular trajectory is coarse-grained into transitions among recurring interaction states.

---

**Fig. S1.**
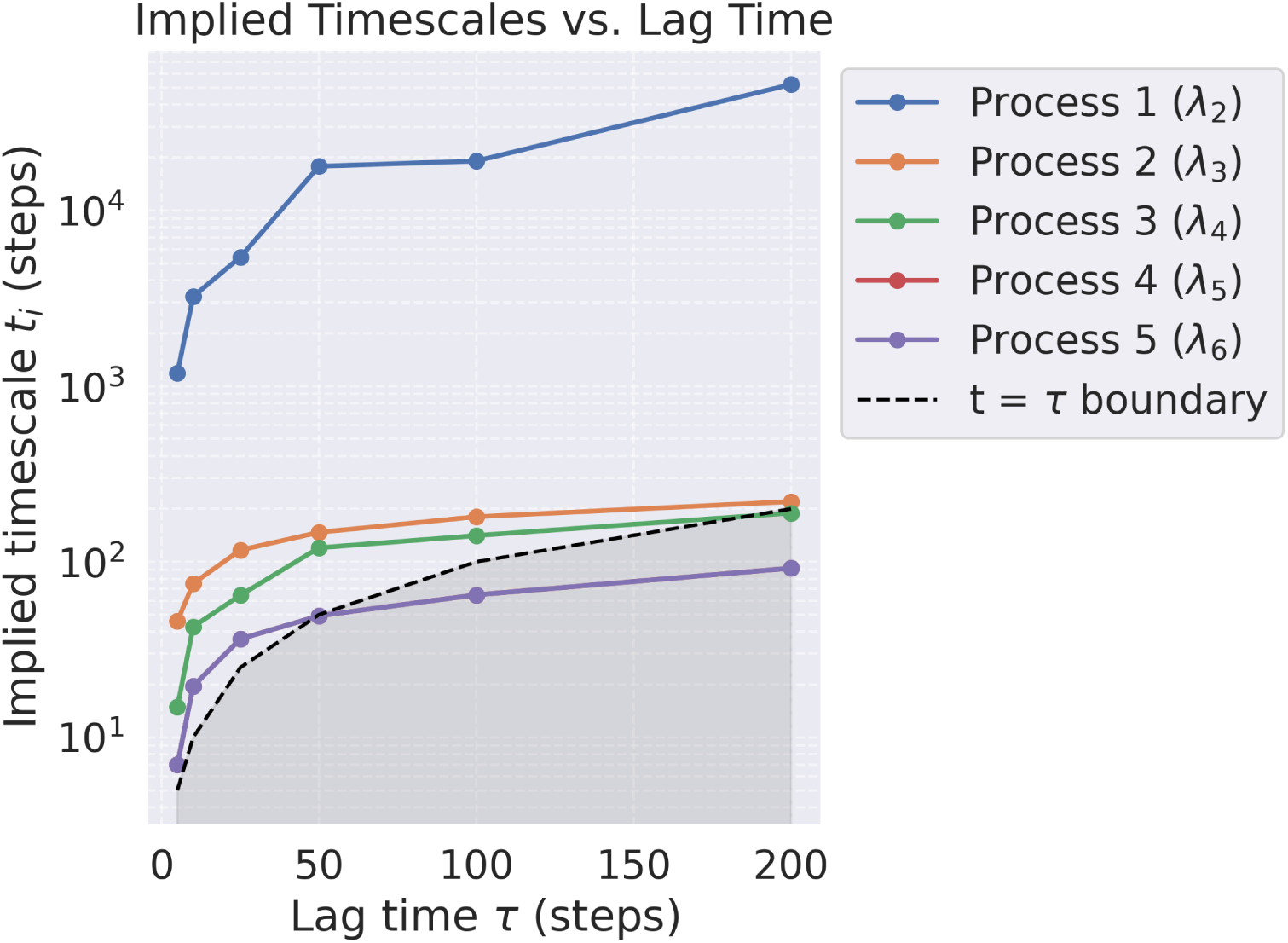
Lag-time selection for the iMSM. Dominant implied timescales are shown as a function of the lag time, expressed in trajectory-sampling steps of 100 ns. The implied timescales become broadly stable near 50 steps (5 *µ*s), which was selected for the subsequent analyses. Unlike a conventional implied-timescale analysis with a fixed state space, the iMSM interaction states were regenerated independently for each candidate lag time because the state construction itself depends on the lag time.

**Fig. S2.**
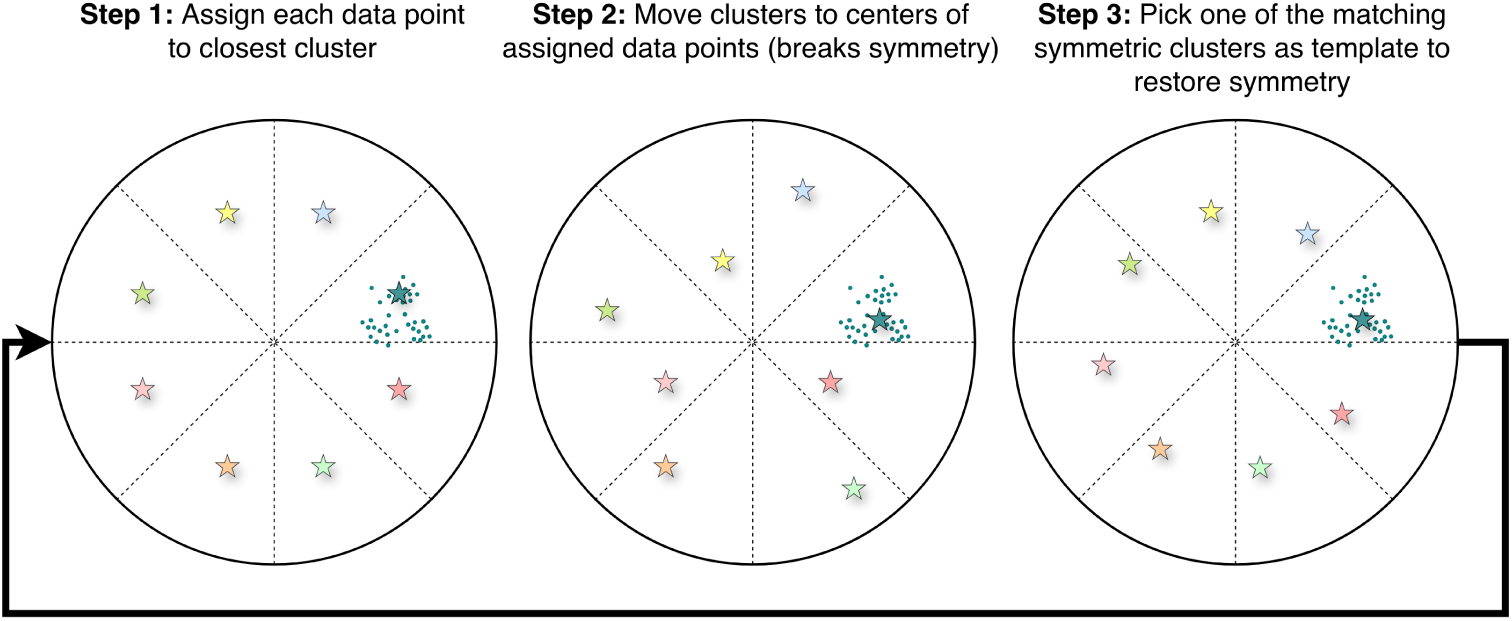
Symmetry-constrained (k)-means clustering used to identify typical interaction states in the nucleocytoplasmic transport system. Each iteration consists of three steps: (1) data points are assigned to the nearest cluster center; (2) each cluster center is updated to the centroid of its assigned points, as in standard (k)-means; and (3) eightfold NPC symmetry is restored in interaction space. Specifically, one center from each symmetry-related set is selected at random, and the remaining seven centers are generated by cyclically remapping interactions with the FG proteins from one NPC spoke to the next. Therefore, remapping does not involve a geometric rotation of the cluster centers.

**Fig. S3.**
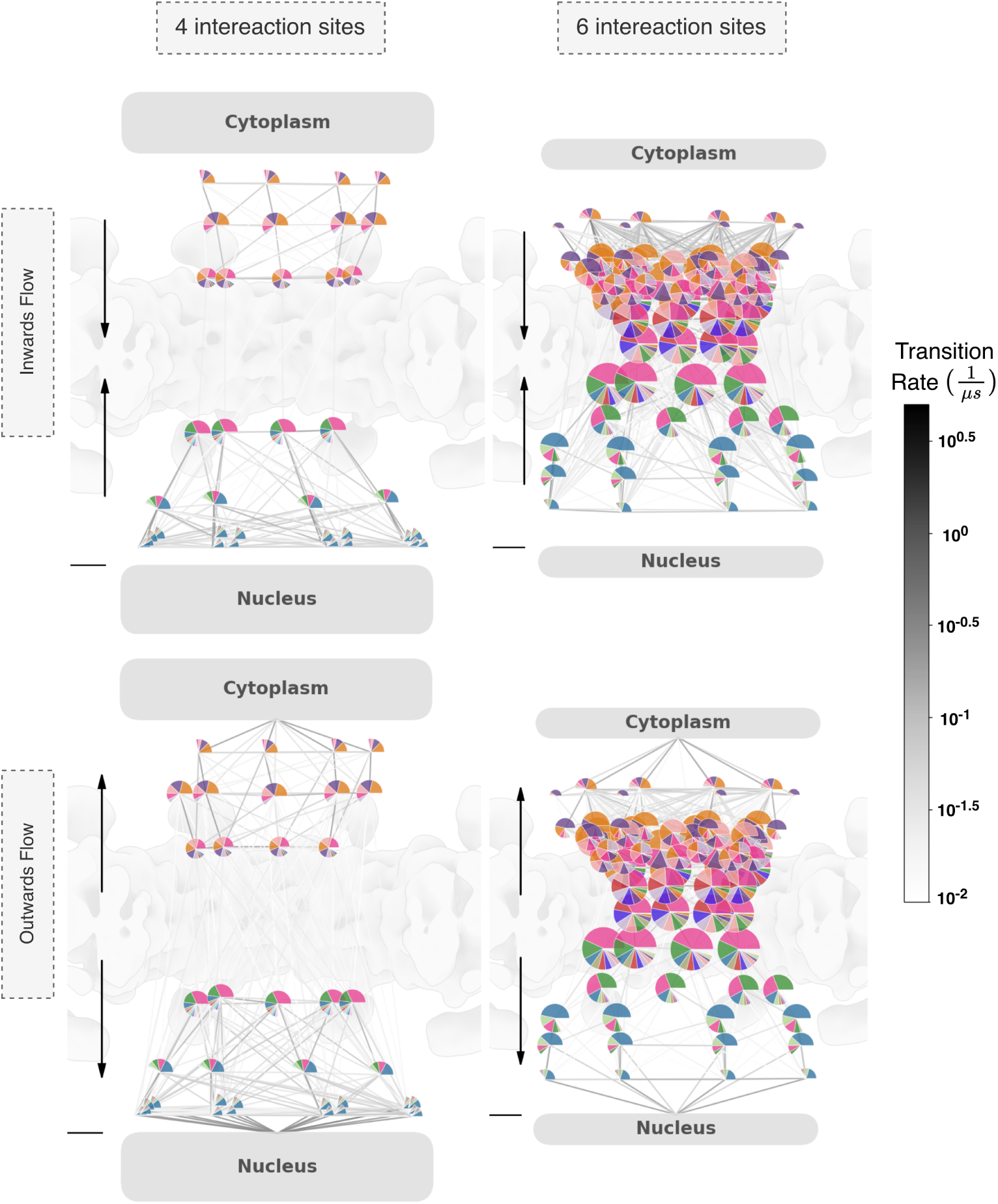
Directional transition rates in representative iMSMs. The iMSMs shown in Fig. 2a for 61-kDa NTRs with four or six FG-Nup interaction sites are reproduced with inward and outward transition rates displayed separately. For each pair of connected states, the inward rate corresponds to the transition from the state with larger *|z|* to the state closer to the pore center (*z* = 0), whereas the outward rate corresponds to the reverse transition. Unlike Fig. 2a, in which each edge represents the average of the two opposing transition rates, each edge here represents a single directional rate. All other graphical elements are as described in Fig. 2a.

**Fig. S4.**
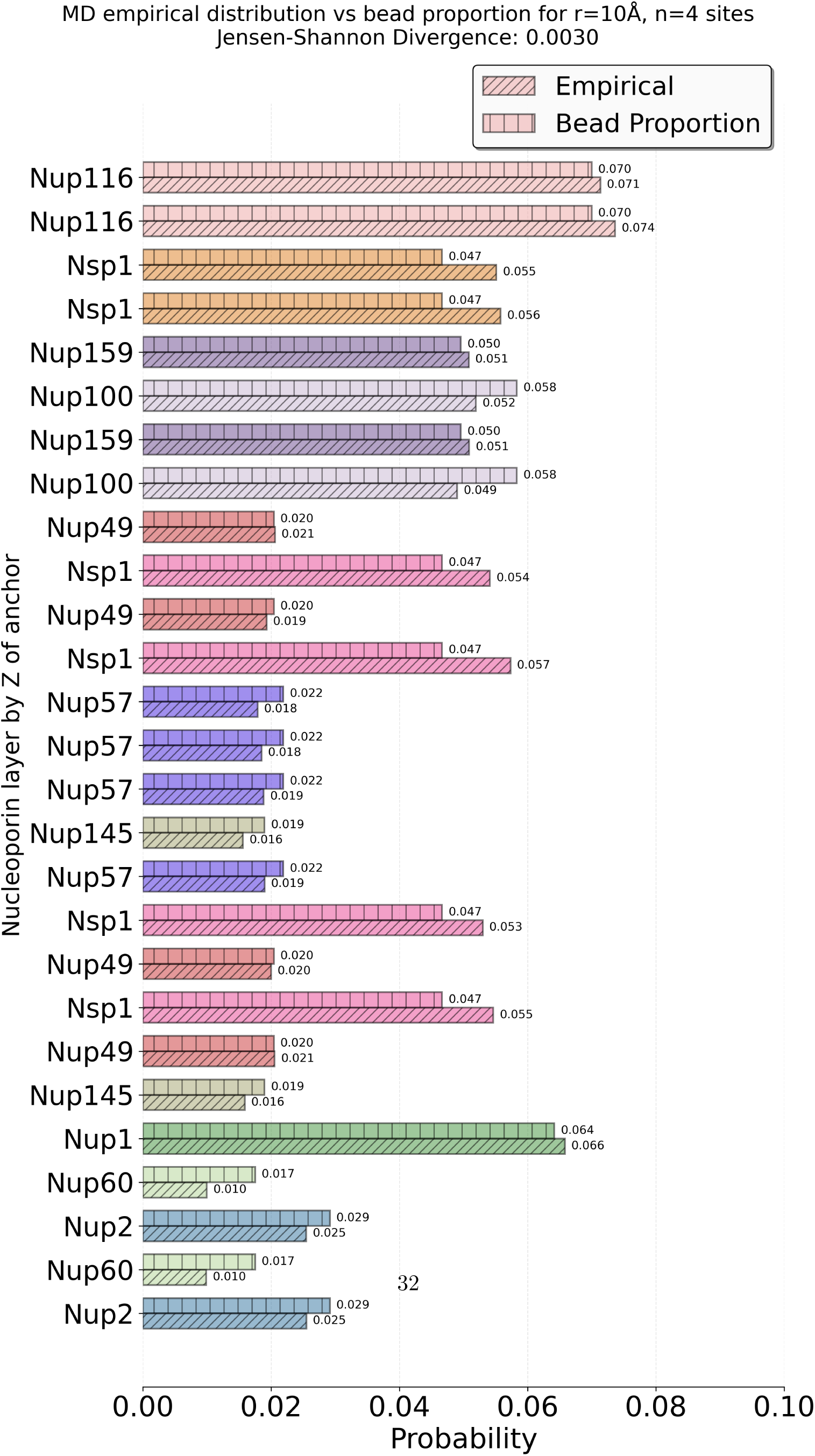
FG-Nup interaction probabilities closely track relative FG-Nup abundance. Empirical interaction probabilities calculated directly from MD trajectories for a 61-kDa NTR with four FG-binding sites in a 54-nm-diameter NPC are compared with the proportion of total FG-Nup beads contributed by each Nup. Nuclear and cytoplasmic unbound states are excluded. The close correspondence indicates that FG-Nup interaction probabilities are largely determined by each Nup’s relative contribution to the FG-repeat mass.

**Fig. S5.**
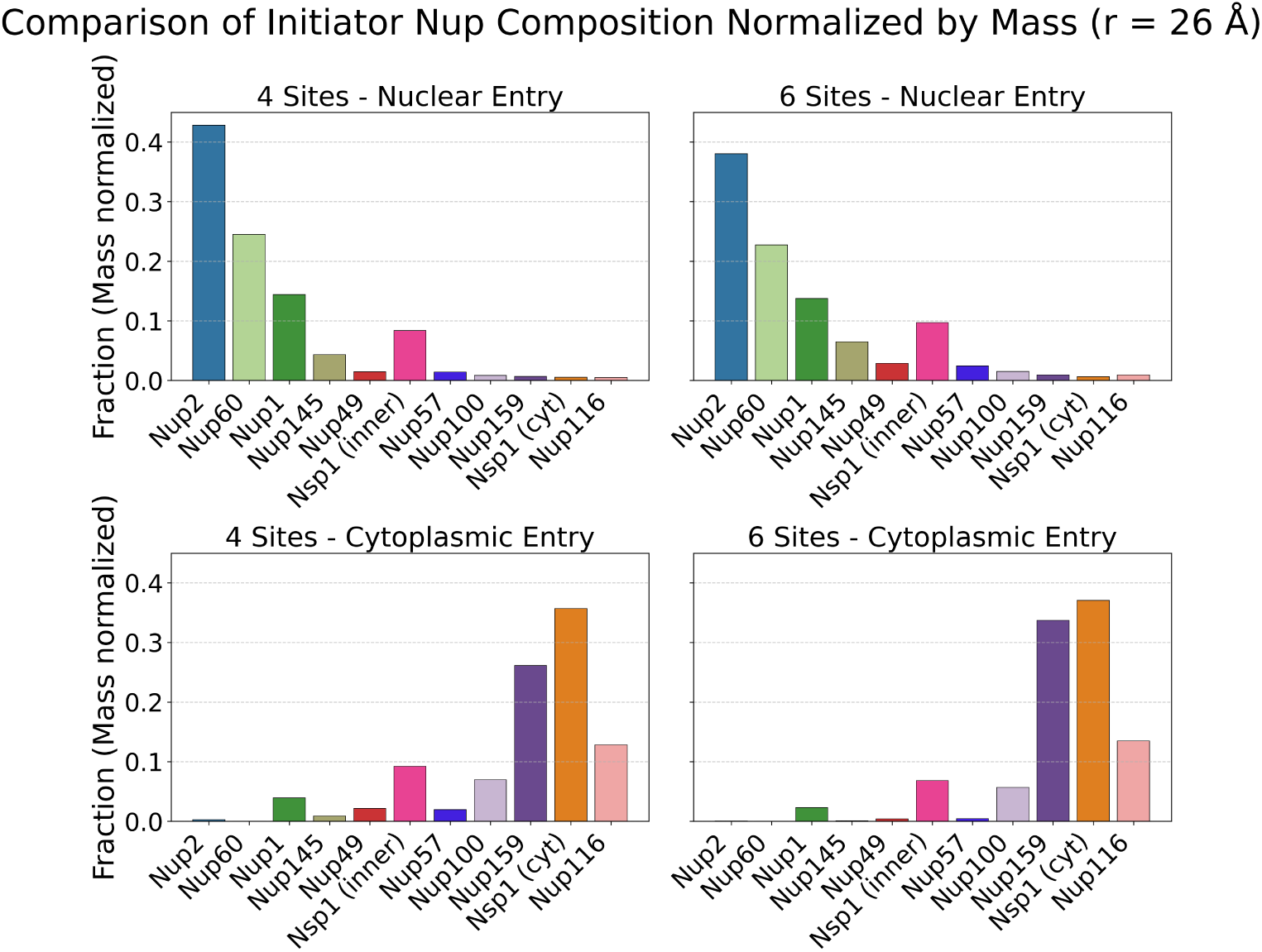
Mass-normalized contributions of FG Nups to transport initiation. Fractions of FG Nups initiating entry from the nuclear side (top) or cytoplasmic side (bottom), calculated from iMSM-generated trajectories for 61-kDa NTRs with four or six FG-binding sites, as in Fig. 2d. For each FG Nup, the initiator fraction was normalized by its modeled FG-repeat mass, calculated as the number of coarse-grained beads per chain multiplied by the number of chains in the NPC. This normalization accounts for differences in FG-Nup chain length and stoichiometry and highlights Nups that contribute disproportionately to initial NTR docking relative to their abundance.

**Fig. S6.**
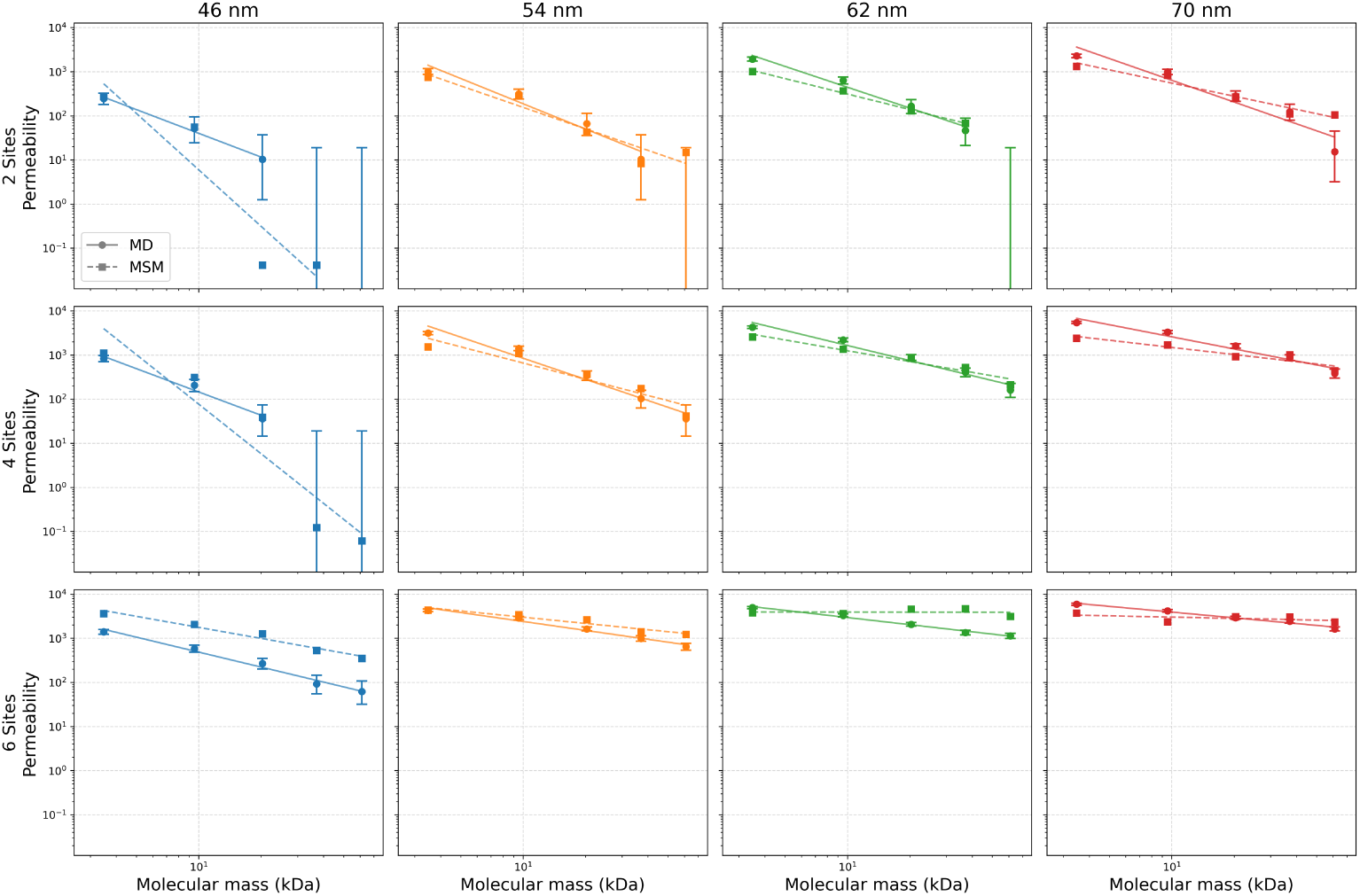
Comparison of empirical MD and iMSM-computed permeabilities across pore variants. As in Fig. 3b, empirical MD permeabilities are shown as solid lines and iMSM-computed permeabilities as dashed lines. Each panel corresponds to a different combination of pore diameter and number of binding sites and shows the permeability of each NTR variant. Missing points indicate conditions for which insufficient statistics precluded a reliable permeability estimate.

**Fig. S7.**
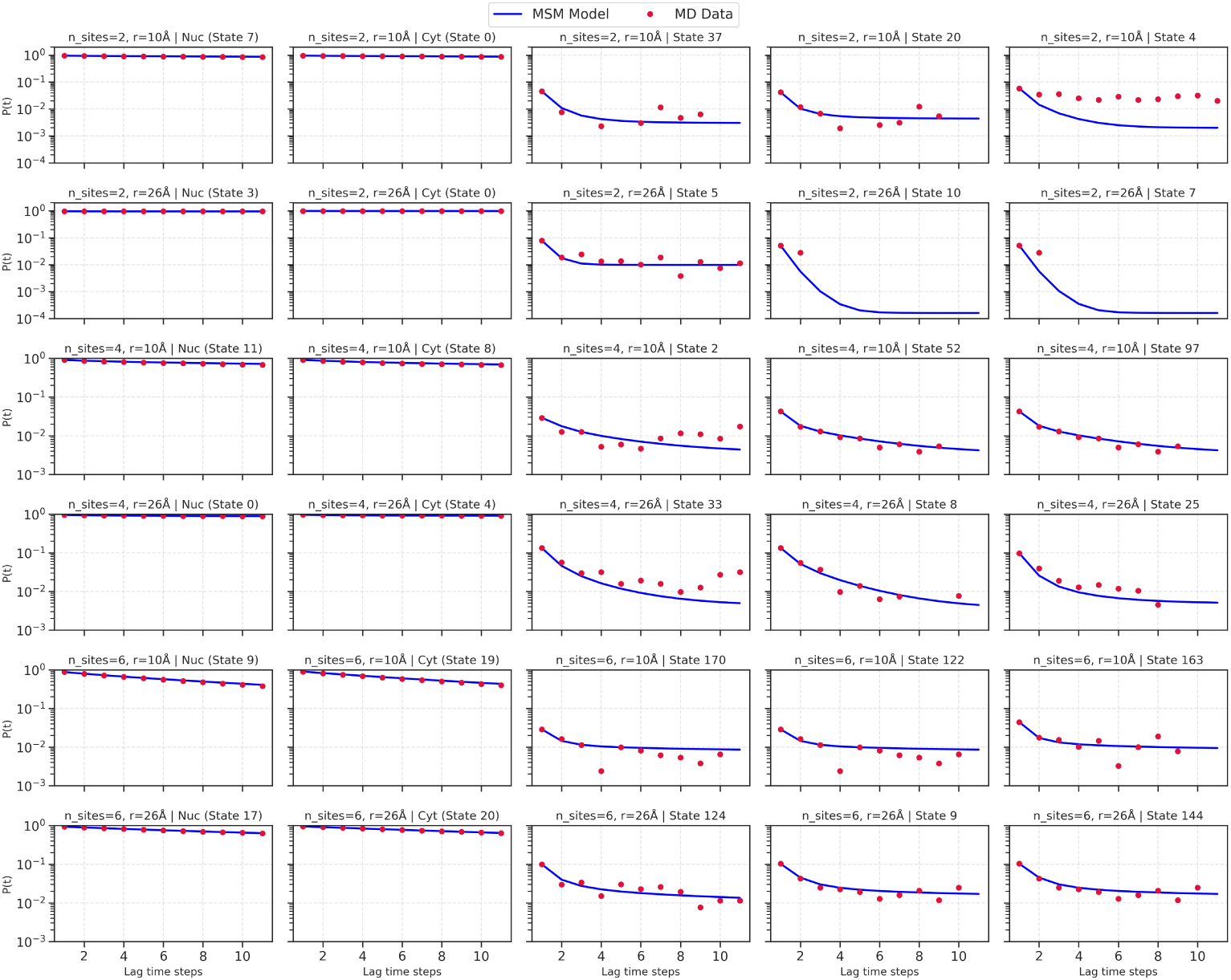
Chapman–Kolmogorov validation of the iMSM models for representative NTR variants. Rows correspond to the smallest and largest NTRs, by molecular weight, with 2, 4, or 6 interaction sites, yielding six NTR variants in total. Columns show five states for each NTR: the nuclear and cytoplasmic states (first two columns) and three randomly selected intermediate states (last three columns). Each panel shows the probability of occupying the indicated state as a function of lag time, comparing estimates obtained directly from the state-assigned molecular dynamics trajectories with predictions from the corresponding iMSM. The overall agreement indicates that the iMSMs reproduce the lag-time-dependent state probabilities observed in the molecular dynamics simulations.

**Fig. S8.**
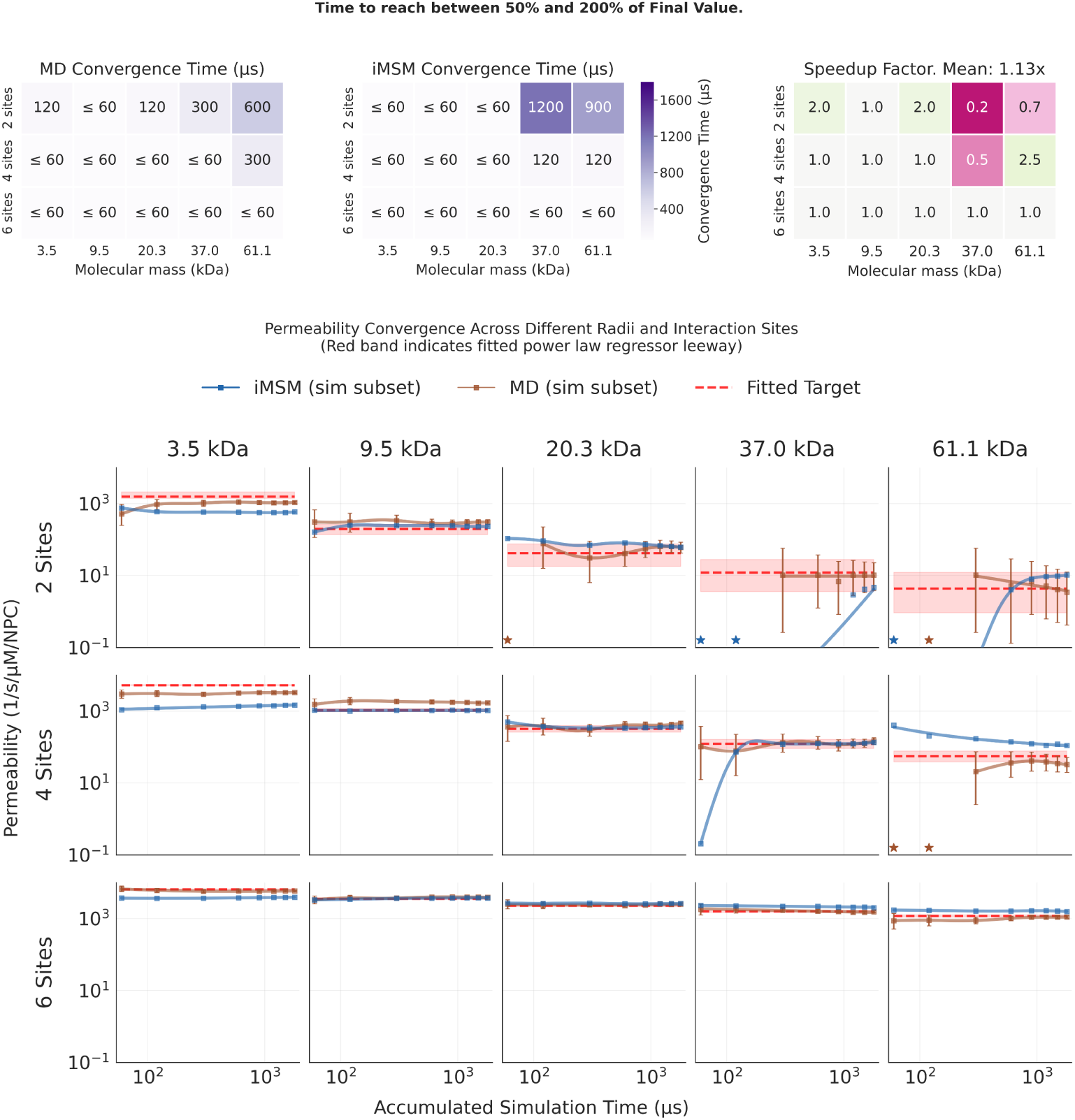
Permeability convergence with increasing numbers of full-length trajectories. **(A)** Convergence times obtained by direct MD transport tallying (left) and iMSMs (center) for NTRs spanning five molecular masses and two, four, or six FG-binding sites. Progressively larger subsets of complete 60 *µ*s trajectories were used, such that a subset containing *n* trajectories had an accumulated simulation time of 60*n µ*s. Values indicate the earliest accumulated simulation time at which the permeability estimate entered and subsequently remained within twofold of its reference value. Corresponding MD-to-iMSM speedup factors are shown at right; the mean speedup across all variants was 1.13-fold. Values of *≤* 60 *µ*s indicate convergence at or before the smallest subset evaluated. **(B)** Permeability estimates from iMSMs (blue) and direct MD tallying (orange) as a function of accumulated simulation time, obtained by progressively increasing the number of complete trajectories included. Red dashed lines show permeabilities obtained from full-data MD power-law fits across molecular masses, fitted separately for each number of FG-binding sites; shaded regions indicate the fit error. Orange error bars show 95% Poisson confidence intervals. Stars indicate subsets with zero observed transport events.

**Fig. S9.**
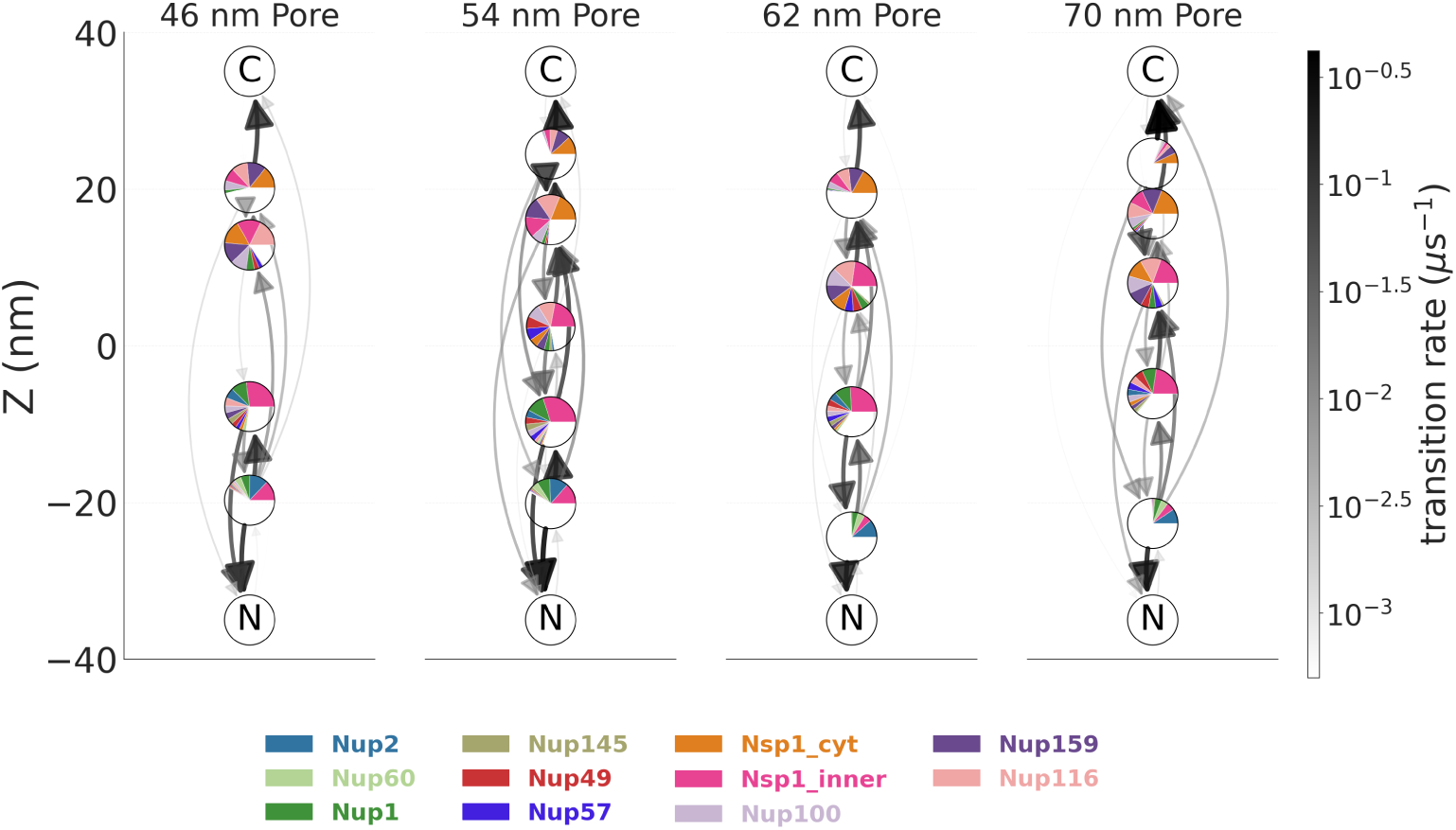
Single-spoke iMSMs colored by FG-Nup type across pore diameters. Single-spoke representations of the iMSMs shown in Fig. 5b–d for a 20.3-kDa NTR with four FG-binding sites across the indicated pore diameters. As in Fig. 2a, colored node wedges indicate the contributions of individual FG-Nup types, whereas white wedges represent unoccupied reference interaction capacity. All other graphical elements are as described in Fig. 5b.

## Notes

https://github.com/ravehlab/iMSM

### Competing Interest Statement

The authors have declared no competing interest.

